# CRY1 couples blue light dependent systemic acquired defense through NPR1 interaction and SnRK3.6-mediated translocation

**DOI:** 10.64898/2026.01.27.701904

**Authors:** Nidhi Singh, Gautam Jamra, Sudip Chattopadhyay, Debasis Chattopadhyay

**Affiliations:** BRIC-National Institute of Plant Genome research, Aruna Asaf Ali Marg, New Delhi, India 110067; Department of Biotechnology, National Institute of Technology, Durgapur, West Bengal, India 713209

## Abstract

Systemic acquired resistance (SAR) is a whole-plant immune response triggered by localized infection. Cryptochromes (CRY) that are blue light receptors, work together with phytochromes (PHY) to regulate light dependent responses in Arabidopsis. Although the effect of light on plant immunity is fairly well studied, the molecular mechanisms remain unclear. In this study, we show that CRY1 is essential for optimal SAR induction under blue light conditions in *Arabidopsis thaliana*. The *cry1* mutants fail to activate SAR under blue light, as evidenced by reduced expression of defense genes, increased susceptibility to *Pseudomonas syringae* (*Pst*DC3000), and impaired stomatal defense. While SAR is intact in both wild type and *cry1* under white light, blue light selectively reveals critical role of CRY1 in SAR. Interestingly, *npr1* mutants activate SAR under blue light, supporting the fact that blue light can trigger SAR through NPR1 independent pathways. Using biochemical and cell-based assays, we show that CRY1 directly interacts with NPR1 in nucleus in a blue-light–enhanced manner. Overexpression of *CRY1* induces NPR1 levels and enhances SAR. Expression of each of them individually partly rescued SAR, but their co-expression fully reinstated it, implying that SAR relies on interaction between NPR1 and CRY1. Stomatal movement measurements further show that CRY1 and NPR1 act synergistically to promote stomatal closure to limit pathogen entry. We also identify SnRK3.6, a blue-light-inducible kinase, as a novel regulator of CRY1. SnRK3.6 physically interacts with and phosphorylates CRY1, facilitating its nuclear translocation. Our results revealed a novel role of CRY1-NPR1 complex formation and regulation of CRY1 by SnRK3.6 in inducing SAR.

## Introduction

Plants defend themselves against pathogens through innate immune system (da Cunha et al., 2006; De Wit, 2007). The defence responses that plants utilise are various cellular process including recognition of pathogen associated molecular patterns (elicitors) or effectors, activation of MAP kinases, induction of pathogen related proteins (PRs), callose deposition or hypersensitive response (HR) culminating into cell death near the infection site (Abramovitch et al., 2006; Jones and Dangl, 2006; Kaur et al., 2022). The cell death responses, ranging from single-cell HRs to necrotic disease lesions triggers the activation of systemic acquired resistance (SAR) in which the entire plant develops heightened resistance to subsequent pathogen attacks following a prior exposure to elicitors from virulent, avirulent pathogens, non-pathogenic organisms or synthetic chemicals like chitosan or salicylic acid (SA) (Klessig et al., 2018). The activation of SAR in the whole plant is associated with activated salicylic acid signaling that induces the expression of defence marker genes through modulating the transcription activator NPR1 (Non-expressor-of-PR1-genes). Upon pathogen challenge, NPR1 is translocated from the cytoplasm to the nucleus, where it interacts with TGA (TGACG-binding) transcription factors to activate the expression of multiple PR genes (Shah and Zeier, 2013; Klessig et al., 2018).

The successful disease progression requires a susceptible plant, a virulent pathogen, and favorable environmental conditions including light (Velásquez et al., 2018). Plants show maximum photosynthetic activity under the wavelength of blue (BL) and red light (RL). The energy of specific wavelength of light depends upon the changing or varying day time, latitude of the area, season, and density of the plant population or shade surrounding the plants in a field (Ballaré et al., 2012; Ballaré and Pierik, 2017). Plants can adapt a change in environmental conditions by altering their physiology in terms of stomatal movement, leaf positioning, and chloroplast production. Plant perceive light through various photoreceptors like phytochrome A (phyA) and phytochrome B (phyB) primarily for red light (Roman et al., 1994; Briggs, 2014), Cryptophromes for blue light (Cashmore et al., 1988), and Ultraviolet 8 receptor (UVr8) for UV-B light. Other receptors like Phototropins (Phot1 and Phot2) (Briggs, 2014) and Kelch-containing F-box protein (KFB) (Christie et al., 2015) are also used by plants to perceive light. The significance of light in host-pathogen interaction is becoming increasingly evident. Diminished responses to various viral, bacterial, and fungal pathogen infection have frequently been observed under dark conditions. For instance, exposure to 8 hrs of light was shown to be required to develop HR after infecting a rice cultivar carrying Xa-10 with an avirulent strain of Xanthomonas oryzae (Guo et al., 1993). According to another report, challenging Arabidopsis with *Pseudomonas syringae* pv. *maculicola* containing *avr* gene in the dark led to increased bacterial growth and reduced local resistance as compared to the infections under light (Katagiri et al., 2002). Availability of prolonged light after infection was shown to be required for enhanced immune response. Induction of SAR response and SA-dependent systemic defence reaction were compromised in red light receptor mutant *phyAphyB*. The *PR1* expression induced by SA and the hypersensitive response (HR) to pathogen infection are reduced in dark or dim light, demonstrating a strict dependence on phyA and phyB. However, the blue-light receptor mutants *cry1cry2* and *phot1phot2* are both capable of establishing a full systemic acquired resistance (SAR) response, suggesting induction of SAR depends on the wave length of light (Griebel and Zeier, 2008). Although, both the SA-induced PR gene expression and HR are reduced in low intensity light, functional chloroplasts was essential for HR. However, this is not required for light-dependent *PR* induction indicating light-dependent HR can occur without induction of SA-mediated defense signaling (Genoud et al., 2002a). Influence of light on plant defense response extends to virus infection also. Resistance to Turnip Crinkle Virus (TCV) infection and HR are dependent on light. A dark treatment immediately after TCV inoculation suppressed HR, resistance and activation of the majority of the TCV-induced genes. However, absence of light did not affect the induction of SA and expression of resistance gene in response to TCV infection. Resistance against TCV is conferred by *HRT* (Hypersensitive Response to TCV) gene. Resistance gene HRT confers resistance to Turnip Crinkle Virus (TCV) in Arabidopsis through a light-dependent pathway that operates independent of the photoreceptors phyA and phyB suggesting light receptors other than phyA and phyB have roles in light-mediated defense response (Chandra-Shekara et al., 2006). Some of the defence responses are independent of light, for example, biosynthesis of Jasmonic acid and camalexin (Zeier et al., 2004). The above reports indicate that light intensity and wave length play an important role in disease progression, with various mechanisms through which light receptors transmit light signals to regulate plant immunity.

Growing body of evidence suggests more contribution of the photoreceptors to plant immunity. Photoreceptor mediated perception of low light intensity modulates plant immune responses (Breen et al., 2023). The inactivation of red-light receptor PhyB generally increases susceptibility to pathogens. In tomato, phyB enhances defense against *Spodoptera eridania*, while both phyA and phyB contribute defence to Arabidopsis against *Pseudomonas syringae* DC3000 (Genoud et al., 2002b; Xiang et al., 2022). RL influences plant defense through JA signaling and JAZ activity, which mediate responses to necrotrophic pathogens and herbivory. PhyB regulates JA biosynthesis genes and enhances JA-dependent defense against *Botrytis cinerea* (Xiang et al., 2022). High R:FR ratios also upregulate SA-mediated signaling, enhancing resistance to biotrophic pathogens. RL exposure increases SA levels, inducing SA signaling and ROS (Reactive oxygen species) production and defense gene activation in a HY5-dependent manner (Gallé et al., 2021). However, its role in RL-induced cell death *via* SA signaling remains unclear. RL receptors PhyA and PhyB contribute to *Pto* DC3000 defense through SA pathways. Low R:FR represses SA defenses by reducing NPR1 phosphorylation and relocalization. phyB mutants show higher *Pto* DC3000 susceptibility, and NPR1 silencing compromises RL-induced resistance indicating a role of NPR1 in the RL-induced resistance. Taken together, RL and FRL regulate plant defense responses in addition to controlling photomorphogenesis, shade avoidance, hormone dynamics, and key transcription activators for environmental adaptation.

In Arabidospsis, NPR1 is considered as master regulator of SAR. As we discussed above, the role of NPR1 in modulating SAR through the expression of PR genes is well established in Arabidopsis. Salicylic acid induces SAR by activating the NPR1 by converting its polymeric form to a monomeric form. This SA mediated signals and SnRK2.8 dependent phosphorylation are required to activate NPR1 and further activation of SAR in Arabidopsis (Lee et al., 2015a). However, the role of NPR1 in SAR induction under varying light intensity and wave lengths and how systemic responses are connected to blue light perception through cryptochrome are not investigated yet.

Regulation of stomatal aperture is a hallmark of plant defense response. Plants regulate stomatal aperture for transpiration, gas exchange and also in response to pathogens. Stomatal aperture is regulated in response to blue light (Singh et al., 2025). Arabidopsis blue light receptor CRY1 mediate pathogen-triggered stomatal closure and, thereby, plant immunity through a light-responsive protein LURP1 (LATE UPREGULATED IN RESPONSE TO HYALOPERONOSPORA PARASITICA) (Hao et al., 2025a). CRY1 enhances R-protein-mediated resistance and PR gene expression against *Pto* DC3000 (Avr*Rpt2*) in Arabidopsis (Liang and Quan, 2010; Wu and Yang, 2010), while CRY2 and phot2 stabilize HRT for TCV resistance (Jeong et al., 2010). However, CRY-interacting bHLH (CIB1) has been shown to inhibit immunity in response to different PAMPs (Malinovsky et al., 2014). Similarly, another class of blue light protein receptor Phot1 and Phot2 negatively impact plant immunity against *Phytopthora infestans* RXLR effector Pi02860 that cause late blight in potato (Naqvi et al., 2022). All these evidences indicate a differential role of different photoreceptors in regulating plant defense signalling.

Systemic acquired resistance (SAR) provides a long lasting broad spectrum resistance in the healthy tissues of plants against a wide range of pathogens. This process primes the uninfected plant tissues and develop preparedness towards future infections. It requires salicylic acid-dependent long-distance mobile signalling and activation of defense response genes in the healthy tissues. (Mou et al.,2003). The evidences mentioned above establish the role of light in modulating plant immunity. The purpose of this study is to investigate the direct molecular connection between SA-mediated signalling and light signalling in SAR. We examined how NPR1 and CRY1 interact in a blue light-dependent manner during systemic acquired resistance (SAR) in *Arabidopsis*. Under low or blue light, *cry1* mutants failed to induce SAR, while *npr1* mutants showed partial induction. *cry1* mutants were more susceptible upon secondary infection and showed reduced *PR1* and *NPR1* expression, as well as more open stomata compared to the wild type plants (Col-0). In contrast, *npr1* mutants had fewer open stomata under blue light. Constitutive expression of either of the genes partially restored SAR, while co-expression of NPR1 and CRY1 fully restored the phenotype of the wild type plants, indicating that SAR depends on both of their presence. We showed that NPR1 and CRY1 interact likely in the nucleus under blue light during secondary *Pseudomonas syringae* infection. NPR1 transcript and protein levels, as well as its nuclear localization, were blue light-dependent. Additionally, SnRK3.6, a kinase involved in blue light perception, interacts with and phosphorylates CRY1, influencing its nuclear localization, similar to how SnRK2.8 phosphorylates NPR1.

## Results

### CRY1 is required for systemic acquired resistance (SAR) under low light conditions

To study the influence of light intensity on the induction of plant defense responses, we first compared the development of SAR in *phyA*, *phyB*, *cry1* and wild type (Col-0) at normal light (100 µmol /m^2^ /s) and low light (40 µmol /m^2^ /s) intensities under 8h-light/16h-dark photoperiod at 22^0^C. The cell death or HR-inducing bacteria *Pseudomonas syringae* pv. *tomato AvrRpt2* (*Pst*AvrRpt2) carrying the *AvrRpt2* avirulent gene was used as primary challenge in the primary tissue (first true leaves) while *P. syringae* pv. *tomato* DC3000 (*Pst*DC3000) was used for secondary challenge in the systemic tissues after 48 hrs from the primary challenge. The mock plants were infiltrated with 10mM MgCl_2_ as the primary challenge while *PstAvrRpt2* was used as the secondary challenge. We observed development of SAR and HR in the blue light receptor *cry1* mutant at normal light condition, wheras the red light receptor mutants phyA and phyB did not show SAR in this condition. However, this observation was altered under low light condition where *cry1* mutant became more sensitive to the secondary challenge in comparison to the mock treated sample. The *phyA* and *phyB* mutants did not show any differential result under normal and low light conditions (Figure 1A-C). It is well known that SAR induction is associated with enhanced accumulation of defense markers like *PR* genes in the distal leaves tissue. We compared the expression of *PR1*, and *NPR1*, the major regulator of SAR, in the systemic tissues 24h post-secondary challenge of *Pst*DC3000 in both mock and SAR samples of *cry1* mutant and wild type (Col-0). Elevated expression of *PR* genes and *NPR1* was observed in *cry1* mutants in comparison to the mock samples under normal light intensity whereas it was comparable to the mock samples under low light growth condition (Figure 1D and 1E), indicating CRY1 is required to activate SAR in the distal leaves under low light condition (Figure 1D-G).

**Figure 1:**
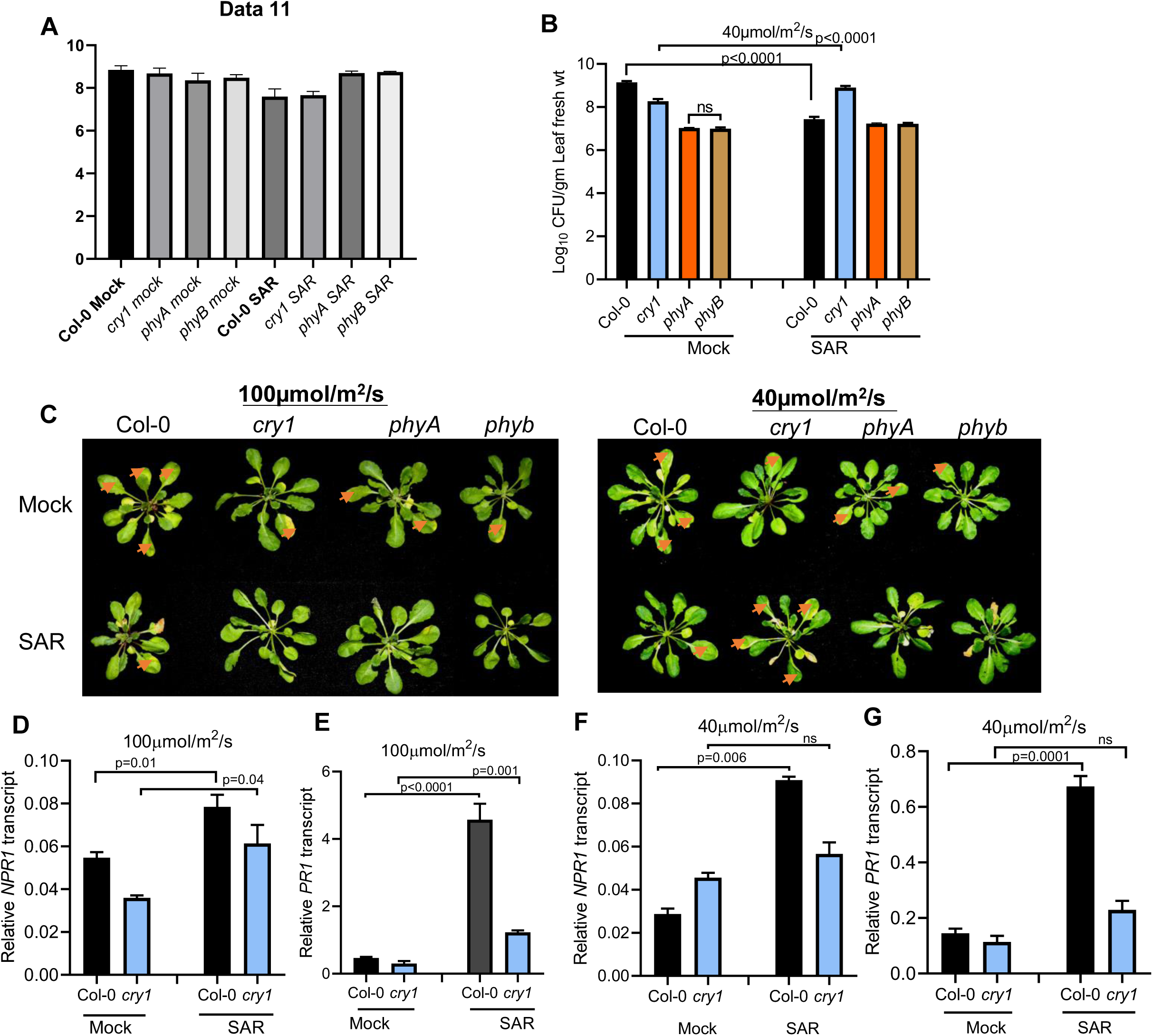
AtCRY1 positively influence systemic acquired resistance (SAR) under low light condition. (A) and (B) *Pseudomonas syringae* pv. *tomato* (*Pst*DC3000) counts in Col-0 (WT), *cry1, phyA, and phyB* mutant plants, growing at two different light intensities. Plants were grown at 100µmol/m^2^/s and 40µmol/m^2^/s light intensities. Bacterial numbers were determined at 3 days post-infiltration (dpi). Each bar indicates the mean standard deviation (SD) (n = 4, independent biological samples), as p values above the compared genotype, as obtained by two-way ANOVA (Tukey’s multiple comparison test). Mock (primary treatment =10 mM MgCl2 and secondary treatment = *Pst*DC3000); SAR (primary treatment = Avr-*Pst* and secondary treatment = *Pst*DC3000). (C) disease phenotype of Col-0, *cry1, phyA,* and *phyB* mutants at 100µmol/m^2^/s and 40µmol/m^2^/s intensity of light (the phenotype of *cry1* for individual leaf is present in supplementary fig 1A). The Avr-*Pst* inoculated primary leaves in the SAR plants show the death of leaves (dried yellow symptoms) (D) and (F) The relative expression of *NPR1* in distal tissues of both Col-0 and *cry1* plants at 24 hours post-inoculation (hpi) with *Pst* DC3000, following secondary challenge. Plants were primarily treated with mock and *Avr* under light intensities of 100 µmol/m^2^/s and 40 µmol/m^2^/s, respectively. (E) and (G) Relative *PR1* expression was quantified in distal tissues of plants treated with mock and Avr following *Pst* DC3000 challenge under 100µmol/m^2^/s and 40µmol/m^2^/s. Each bar indicates the mean standard deviation (SD) (n = 3 independent biological experiment) the significance has been denoted as p values above the compared genotype, as obtained by two-way ANOVA (Tukey’s multiple comparison test). The experiments repeated twice with similar results.

### CRY1 is not required for SAR towards *Pst*DC3000 under low light at high ambient temperature

In nature, changes in light intensity usually come with changes in temperature. To examine the impact of increased ambient temperature on the development of SAR phenotype of *cry1* mutants, we first grew the wild type plant (Col-0) and *cry1* at low light intensity (40 µmol /m^2^/s) and then kept one batch of both the genotypes at 26^0^C and the other at 22^0^C as before under the same light intensities for 36h before giving the primary challenge followed by secondary inoculation. The bacterial load was determined at 3dpi of *Pst*DC3000. Interestingly, *cry1* mutants showed significantly lesser bacterial load as compared to its mock inoculated plants (Figure 2A and 2B) when grown at 26^0^C under low light. The SAR (Figure 2C) phenotype was associated with enhanced expression of defense genes like *PR1* and *NPR1* in the systemic tissues at 26^0^C (Figure 2F-G) while they were not induced, rather suppressed in the systemic tissue of the plant grown at 22^0^C and 40 µmol /m^2^/s light intensity (Figure 2D-E,). These findings suggest that activity of CRY1 with regards to SAR changes with ambient temperature and CRY1 is not required for SAR development at higher temperature, highlighting the interplay between light, temperature, and immunity, and a role of CRY1 in this cross talk.

**Figure 2.**
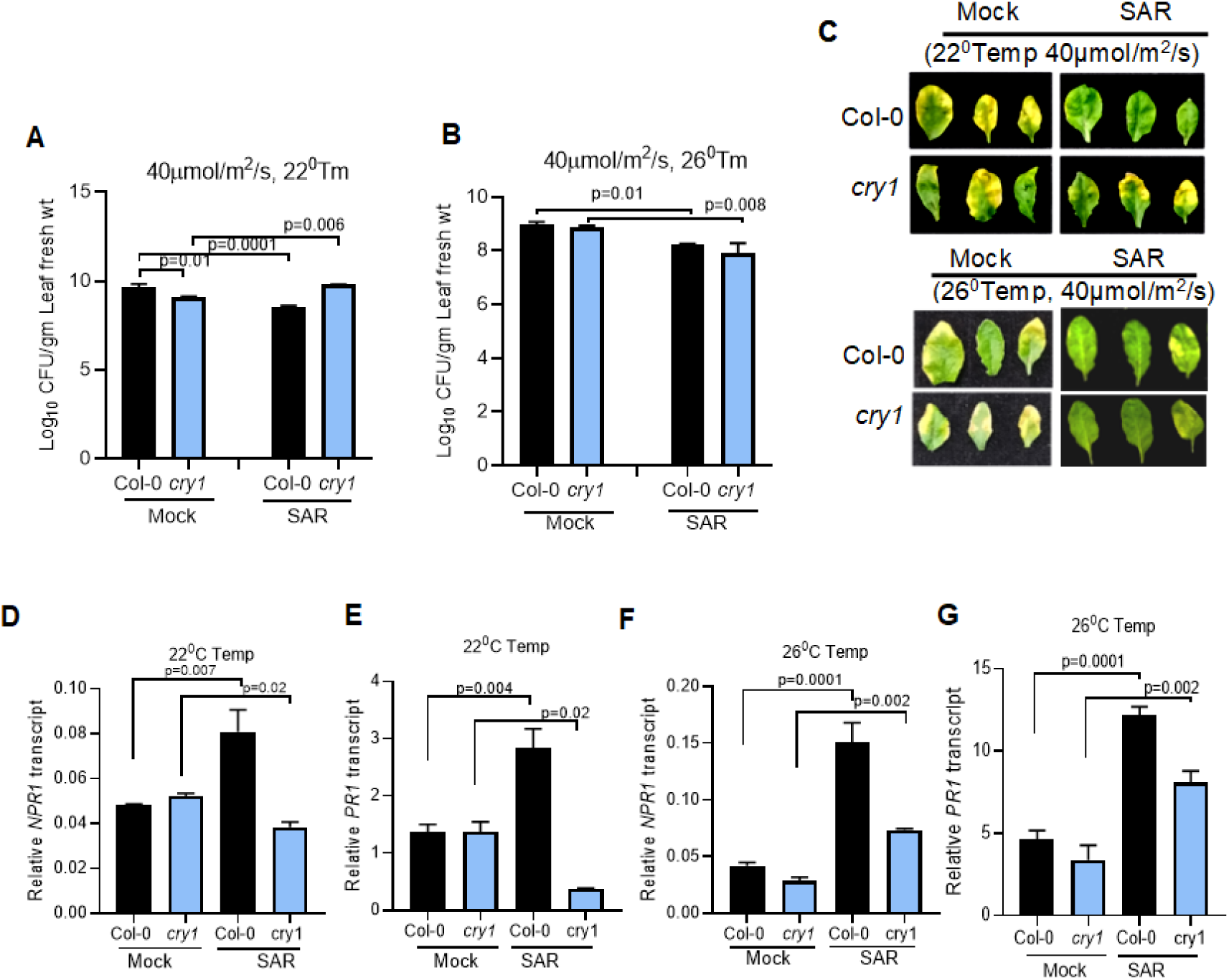
The SAR phenotype of cry1 mutants was restored at higher ambient growth temperature. (A) (A) and (B) *Pseudomonas syringae* pv. *tomato* (*Pst*DC3000) counts in Col-0 (WT) and *cry1* mutant plants, growing at 40µmol/m^2^/s light intensity. Bacterial numbers were determined at 3 days post-infiltration (dpi). Plant were kept at 22^0^C and 26^0^C under 40µ*mol/m^2^/s*, 4 days before primary inoculation and they were kept there until the CFU count test. Mock (primary treatment =10 mM MgCl2 and secondary treatment = *Pst*DC3000); SAR (primary treatment = Avr-*Pst* and secondary treatment = *Pst*DC3000). Each bar indicates the mean standard deviation (SD) (n = 4 independent biological samples) Star/s above the bars indicate significant difference (P < 0.05, 0.001, 0.0001) as obtained by two-way ANOVA (Tukey’s multiple comparison test) (Mock and SAR). Mock (primary treatment =10 mM MgCl2 and secondary treatment = *Pst*DC3000); SAR (primary treatment = Avr-*Pst* and secondary treatment = *Pst*DC3000). (C) disease phenotype of Col-0 and *cry1* mutants at 22^0^C and 26^0^C temperature under 40µmol/m^2^/s intensity of light (D) and (F) The relative expression of *NPR1* in distal tissues of both Col-0 and *cry1* plants at 24 hours post-inoculation (hpi) with *Pst* DC3000, following secondary challenge. Plants kept at either 22^0^C or 26^0^C, were primarily treated with mock and *Avr*, respectively. (E) and (G) Relative *PR1* expression was quantified in distal tissues of plants treated with mock and *Avr* following *Pst* DC3000 challenge. Each bar indicates the mean standard deviation (SD) (n = 3 independent biological experiments) the significance has been denoted as p values above the compared genotype, as obtained by two-way ANOVA (Tukey’s multiple comparison test). The experiments repeated two times with similar results.

### NPR1 and CRY1 may interconnect blue light-mediated systemic acquired resistance and stomatal regulation

To further explore the effect of wave length of the imposed light on the development of systemic acquired resistance (SAR), we compared the growth of *Pst*DC3000 in the systemic leaf tissue of the wild type (Col-0) and *cry1* mutants under white light and blue light conditions, since light spectrum often influence certain biological processes similarly (Liu and van Iersel, 2021; Wu et al., 2024). We also included *npr1* mutant in this experiment as it is known to be insensitive after secondary challenge under normal light and growth condition (Backer et al., 2019a; Zavaliev and Dong, 2024). The systemic leaf tissue of *cry1* mutant exhibited a heightened sensitivity compared to the wild type plants (Figure 3B and 3C) under blue light which was associated with reduced accumulation of defense-related genes such as *PR1* in comparison to the wild type (Col-0) plants (Figure 3D). The above results highlight the role of cryptochrome in blue light-dependent defense regulation.

**Figure 3:**
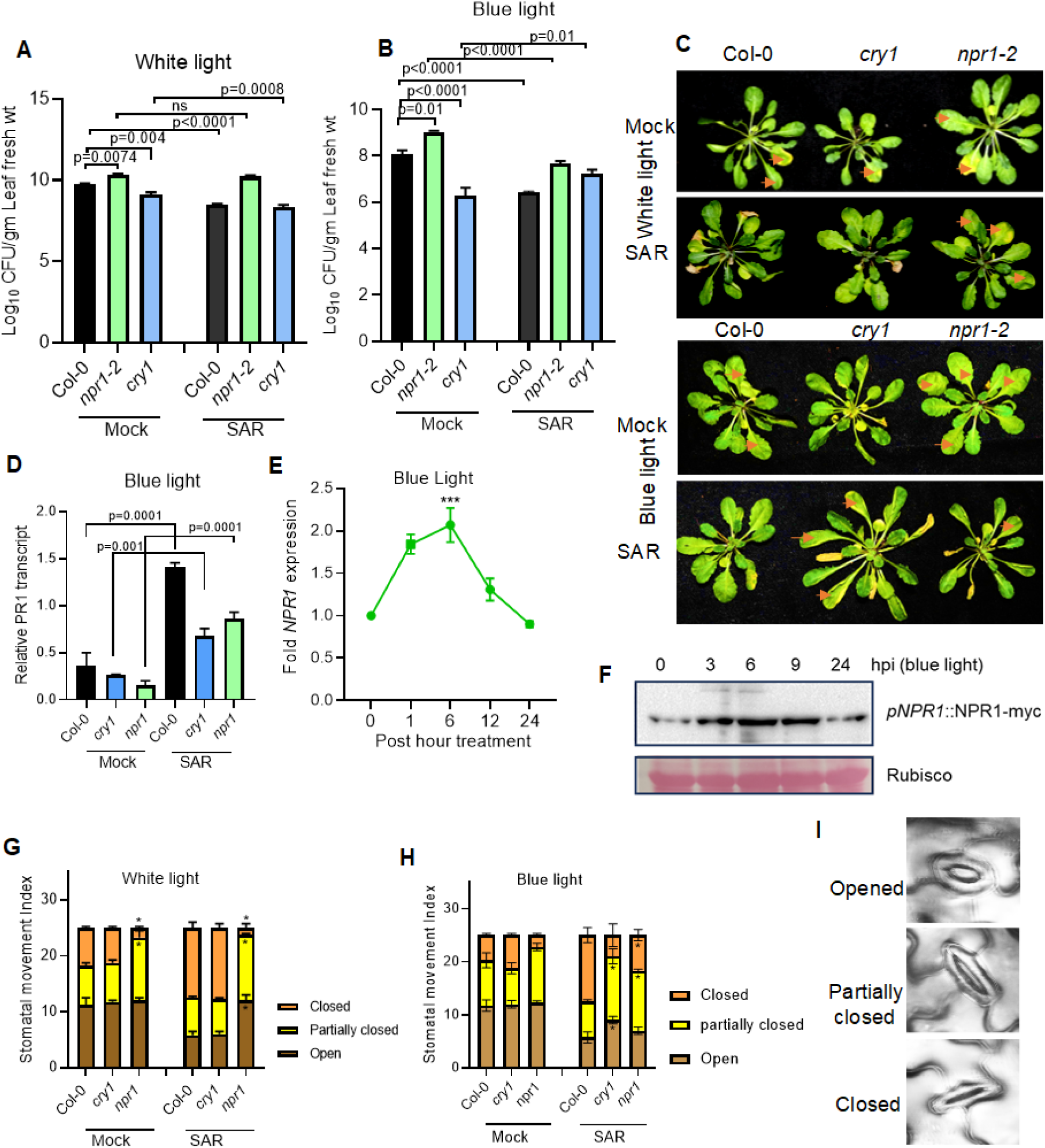
Function of CRY1 and NPR1 in SAR and stomatal movement under blue light. (A and B) Bacterial count in systemic tissue under white and blue light conditions. Plant were kept under blue light two days before primary mock and *Avr* inoculation and they were kept there until the CFU count test after secondary challenge with *Pst* DC3000. (C) disease phenotype of the indicated genotypes under blue light after secondary challenge with *Pst*DC3000. The brown arrows show the disease symptoms on the leaves, the lower died leaves show the primary inoculation with *Pst*-Avr in SAR. Each bar indicates the mean standard deviation (SD) (n = 4 independent biological samples) as obtained by two-way ANOVA (Tukey’s multiple comparison test) (D) Expression of *PR1* under white (supplementary figure 2B) and blue light in the systemic tissue 24 post hour secondary inoculation, (E) fold expression of *NPR1* at 6hpi is significantly high as compared to 0hpi (significance was tested through one-way ANOVA, (number of samples = 3 from three independent biological samples) where **, p≤ 0.001, ***. p≤ 0.0001. (F) NPR1 protein accumulation over the time course of indicated time points under blue light in Col-0 *pNPR1::NPR1-cmyc* plants respectively. leaves tissue was harvested at the indicated time point. After extracting total protein, immunoblot was developed using anti-myc antibody (G-I) Stomatal movement index was quantified under blue and white light. After 48 hours of primary inoculation with mock and *Avr,* systemic leaves were detached and kept in stomatal opening solution to completely opened the stomata for 2 hours, then kept in 5uM *flg22* solution. After 3-hrs of flg22 treatment stomata were observed under confocal microscope. Stomata (n=50) were counted and categorised as open, partially opened, and closed (shown in G), from three independent biological samples.

Since *cry1* mutant exhibited reduced *NPR1* expression in systemic tissues under low light and blue light conditions (Figure. 2D), leading to lower levels of *PR1*, and *npr1* mutants displayed a SAR-insensitive phenotype under normal light conditions, we aimed to explore the connection between NPR1 and CRY1. To investigate this, we first analyzed NPR1 expression under blue light treatment over a 24-hour period. Both *NPR1* transcript and protein levels showed induction in a blue light-dependent manner (Figure 3E and 3F), as *CRY1* (Supplementary Figure 1A). We aimed to investigate the role of NPR1 in systemic acquired resistance (SAR) under blue light. To do so, we assessed the *Pst*DC3000 load in the systemic tissues of *npr1-2* mutants following a primary challenge with *Pst*AvrRpt2 in the primary leaves under blue light. Interestingly, the *npr1-2* mutant exhibited a SAR phenotype under blue light conditions (Figure 3B, 3C, and Supplementary Figure 1E), unlike in normal light intensities (Figure 3A, 3C, and Supplementary Figure 1B and 1C), as previously reported (Cao et al., 1997). The SAR phenotype under blue light was also accompanied with enhanced *PR1* gene expression as compared to mock treated systemic tissues at 24dpi (Figure 3D). The blue light–induced SAR phenotype with higher *PR1* expression may indicate that NPR1-independent pathways also contribute to blue light-dependent SAR signaling (Uquillas et al., 2004; Blanco et al., 2005; Singh et al., 2018; Backer et al., 2019b).

Previous studies have illustrated that NPR1 plays a crucial role in conferring systemic resistance in plants through stomatal defense following priming with an avirulent pathogen (Guan et al., 2023). To explore the link between blue light and stomatal status in *cry1* and *npr1-2* mutants in SAR, we compared the stomatal index in the systemic tissues of Col-0, *npr1-2*, and *cry1* mutants under white and blue light in mock and SAR leaves tissue. The stomatal index assay was performed on systemic leaves following treatment with *flg22*, after which the leaves were incubated in a stomatal opening solution for 2-3 hours and observed under a confocal microscope. Under white light, no significant differences were observed in the proportion of open, partially closed, and closed stomata between mock-treated and *Avr*-treated systemic tissues of wild type (Col-0) and *cry1* mutants (Figure 3G). As expected, *npr1-2* mutants exhibited a significantly higher number of partially closed stomata in both treatments (Mock and SAR), which may render them more sensitive to *flg22* challenge in systemic leaves (Figure 3G). Additionally, the number of closed stomata was higher and comparable between wild type (Col-0) and *cry1* mutants in systemic tissues (Figure 3G). Under blue light, the stomatal response in mock-treated systemic tissue was similar to that observed under white light (figure 3H). However, *cry1* mutants showed a significantly greater number of partially closed and open stomata compared to wild type (Col-0), whereas *npr1-2* mutants displayed a significantly higher number of partially closed and completely closed stomata as compared to the *cry1* mutants. The stomatal behaviour in *npr1-2* mutants under blue light might have contributed to their observed SAR phenotype (Figure 3H, I), whereas the stomatal behaviour of *cry1* made them unable to induce SAR. These results highlight a potential link between blue light signaling, NPR1 function and CRY1, in regulating stomatal aperture in systemic immunity.

### NPR1 and CRY1 interacts in the nucleus during SAR development and their interaction is blue light dependent

To further investigate the molecular connection between NPR1 and CRY1 in blue light-mediated systemic acquired resistance, we first examined their interaction by *in vitro* GST pull-down assay. Recombinant NPR1 fused with GST and CRY1 fused with His were incubated with GST-Sepharose beads for 6 hours. The pulled-down proteins were then analysed via immunoblotting using anti-GST and anti-His antibodies. The immunoblot results clearly displayed interaction between NPR1 and CRY1, with the input samples confirming the presence of both proteins (Figure 4A).

**Figure 4:**
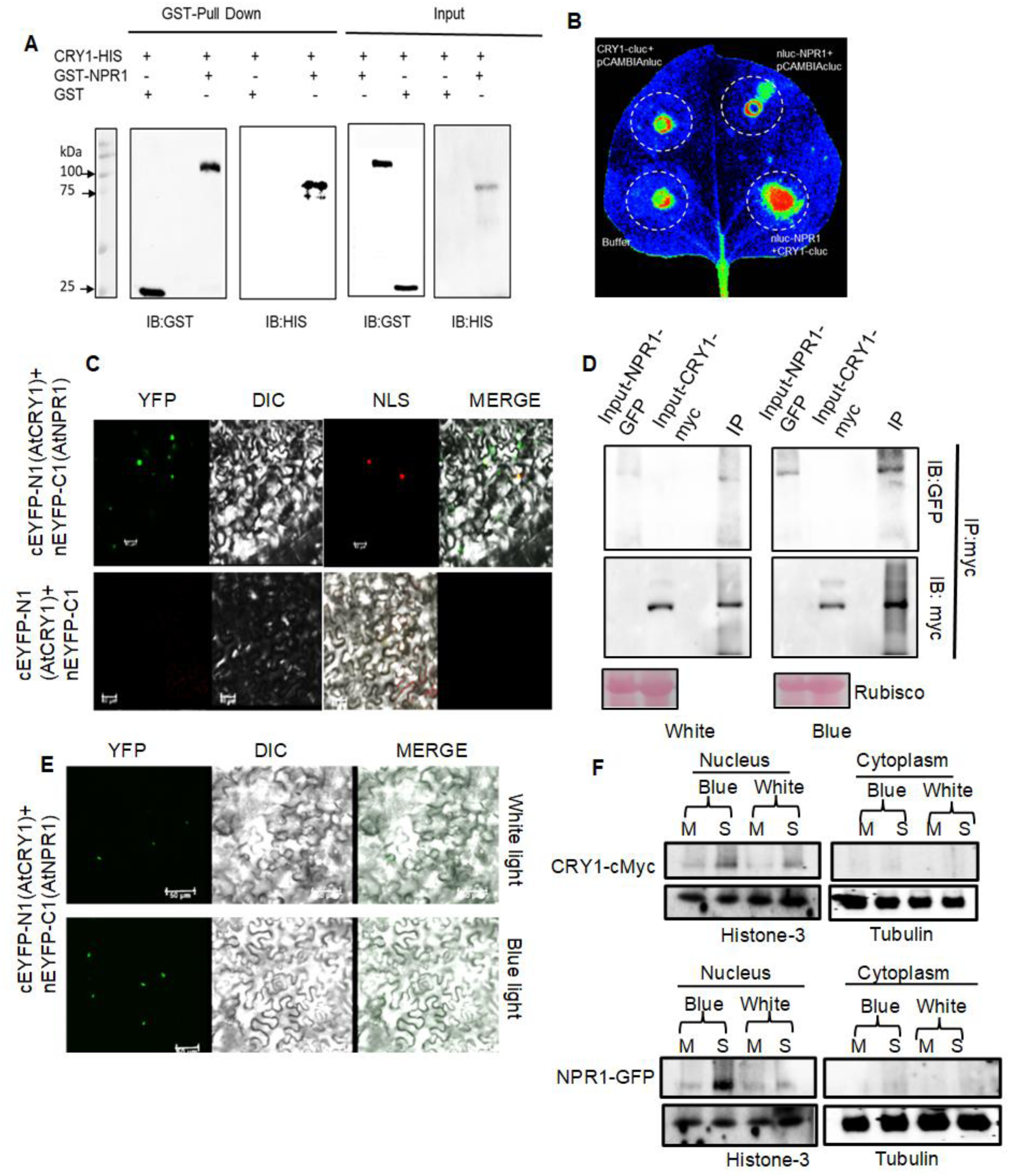
NPR1 and CRY1 Interacts in the nucleus during SAR under blue light treatment. (A) *In vitro*-GST-pull down assay to check the *in vitro* interaction of NPR1 and CRY1. (B) Split luciferase assay to check the invivo interaction between NPR1 and CRY1 in *N. benthamiana* leaves. (C) BiFC assay to show the *in vivo subcellular interaction of* NPR1 *and* CRY1 in *Nicotiana* leaves. Both the constructs (cEYFP-N1-AtCRY1 and nEYFP-C1-AtNPR1) were transiently expressed in *N. benthamiana* leaves using *A. tumefaciens* and observed under confocal microscope after 48hrs of infiltration in leaves. (D) CoIP, to show the *In vivo* interaction in *Arabidopsis* plant expressing both *35S::NPR1-GFP* and *35S::CRY1-cmyc* under blue light. Total protein was immunoprecipitated through cmyc antibody followed by immunoblot assay through GFP and cmyc antibody. Equal amount (500 µgm) of total protein was taken for the immunoprecipitation for both the treatment, white and blue light (shown by rubisco). (E) BIFC assay under white and blue light. Just after infiltrating the agrobacterium harboring the NPR1 and CRY1 construct one batch of plants was kept in white light and another in blue light. Confocal imaging was done after 48 hrs of infiltration. (F) Subcellular fractionation assay in the Arabidopsis, expressing both NPR1-GFP and CRY1-cmyc. Light treatment and *Pst*DC3000 inoculation was done as described in the earlier experiments. The equal amount of protein from nuclear (500 µgm) and cytoplasmic (500µgm) fraction was taken for IP with cmyc antibody followed by western blot with both GFP and cmyc antibody to detect and compare the differential accumulation of CRY1 and NPR1 in nucleus and cytoplasm under white and blue light. M and S stand for Mock and SAR.

The interaction was further validated *in vivo* using a split luciferase complementation assay. NPR1 was cloned into pCAMBIA1300-nLUC to generate an NPR1-nLUC fusion, while CRY1 was cloned into pCAMBIA1300-cLUC to create a CRY1-cLUC fusion. Both constructs were transiently co-expressed in *Nicotiana benthamiana*, and their interaction was assessed by the reconstitution of luciferase activity. Luminescence was visually detected after luciferin application at the infiltrated leaf areas. As a negative control, co-infiltration of pCAMBIA1300-nLUC with pCAMBIA1300-cLUC-CRY1 did not restore luciferase activity, confirming the specificity of the NPR1-CRY1 interaction (Figure 4B).

The subcellular localisation of CRY1 and NPR1 in Nicotiana benthamiana leaves is in the plasma membrane (Supplementary Figure 2A) and nucleus (Supplementar Figure 2B) respectively.To know the subcellular localisation of this interaction between NPR1 and CRY1 we performed bimolecular fluorescent complementation assay through co-infiltration of pSITE3CA-NPR1-eYFPn and pSITE3CA-CRY1-eYFPc in *Nicotiana benthamiana*. The reconstitution of YFP fluorescence in the nucleus suggest that the interaction between NPR1 and CRY1 occurs predominantly in the nucleus under blue light (Figure 4C and 4E). Furthermore, to validate the interaction between the major regulator of systemic acquired resistance NPR1 and blue light receptor CRY1, we generated the *35S::NPR1-GFP* (Supplementary Figure 3B) and *35S::CRY1-cmyc* (Supplementary Figure 3A) Arabidopsis plants. Stable homozygous lines were crossed to generate the *35S::NPR1-GFP/35S::CRY1-cmyc* in a single homozygous line (Supplementary Figure 3C). Co-immunoprecipitation was performed to validate *in planta* interaction between NPR1 and CRY1 and its significance under blue and white lights. We observed that the extent of interaction was more in the blue light condition as compared to white light, using an equal amount of protein for immunoprecipitation using c-myc antibody (Figure 4D). In conclusion, our findings provide a strong evidence for a more prevalent interaction between NPR1 and CRY1 in a blue light-dependent manner. Also, these results suggest a crucial role of CRY1 in modulating NPR1 function under blue light, providing new insights into the molecular mechanisms linking blue light signalling and SAR in Arabidopsis. We further validated the subcellular specificity of NPR1 and CRY1 interaction following biochemical method by subcellular fractionation of proteins from the cytoplasm from the local and distal tissues after priming (mock and *Avr* treated). The abundance of NPR1 and CRY1 in the immunoprecipitated protein fractions has been detected through immunoblot (GFP-Ab for NPR1 and c-myc Ab for CRY1) (Figure 4F). We observed that the abundance of NPR1 and CRY1 protein was higher in the nuclear fraction of the systemic tissue under blue light as compared to that in white light. However, NPR1 was accumulated more in the nuclear fraction of the systemic tissue under blue light as compared to CRY1 (Figure 4F). The enhanced nuclear accumulation of both NPR1 and CRY1 under blue light suggested a synergistic interaction between these two proteins in the regulation of SAR. The preferential nuclear localization of NPR1 over CRY1 may indicate an essential role for NPR1 in initiating the transcriptional responses required for SAR, with CRY1 potentially modulating these responses through its light-dependent activities.

### CRY1 Enhances NPR1-Mediated Systemic Acquired Resistance Under Blue Light

To investigate the functional significance of interaction between NPR1 and CRY1 in systemic acquired resistance under blue light, we compared the bacterial load in the systemic tissue in *NPR1* overexpressing (*35S::NPR1-GFP)*, *CRY1* overexpressing (*35S::CRY1-cmyc*), *NPR1*/*CRY1* overexpressing (*35S::NPR1-GFP/35S::CRY1-cmyc*) plants and compared with that in the wild type (Col-0) plants under white and blue light. All genotypes exhibited resistance to the secondary challenge under both white (Supplementary Figure 4A) and blue light (Figure 5A). However, *35S::CRY1-cmyc* plants showed significantly stronger SAR measured by bacterial load as compared to wild type (Col-0) under blue light. The *35S::NPR1-GFP* plants displayed systemic resistance in comparison to wild type Col-0 (Figure 5A). Overexpression of *NPR1* led to *PR1* induction in both local and systemic tissues under both light conditions (Supplementary Figure 4B), and CRY1 overexpression also triggered NPR1 expression in systemic tissue under blue light. However, the double overexpressor *35S::NPR1-GFP*/*35S::CRY1-myc* showed notably much higher SAR in *Avr*-inoculated plants in blue light than both the single overexpressor accompanied by the highest *PR* transcript levels (Figure 5A-C). Analysis of stomatal behavior in systemic leaves (inoculated with 10 mM MgCl₂ or *Pst*AvrRpt2 under blue light) revealed that *NPR1* overexpression increased the number of partially closed stomata, while *CRY1* overexpression further increased both partially and fully closed stomata as compared to wild type (Col-0) (Figure 5D). The *NPR1*+*CRY1* overexpressing plants had the highest proportion of completely closed stomata (Figure 5D and Supplemenatary Figure 4D)), correlating with a significantly reduced *AHA1*(H^+^-ATPase) gene expression, required for the stomatal opening, transcript levels (Figure 5E). The corresponding behaviour under white light is provided in the Supplementary Figure 4C. Altogether, these results support a hypothesis where CRY1, when activated by blue light, potentiates the function of NPR1 leading to stronger *PR1* expression, stomatal defense, and SAR. Also, NPR1 and CRY1 may act synergistically in providing systemic resistance under blue light. Our finding supports the previous reports where CRY1 enhances SA- and light-mediated defense signaling, promoting PR1 expression under blue light (Griebel and Zeier; 2008; Liang and Hong-Quan, 2010; Wu and Yang, 2010).

**Figure 5:**
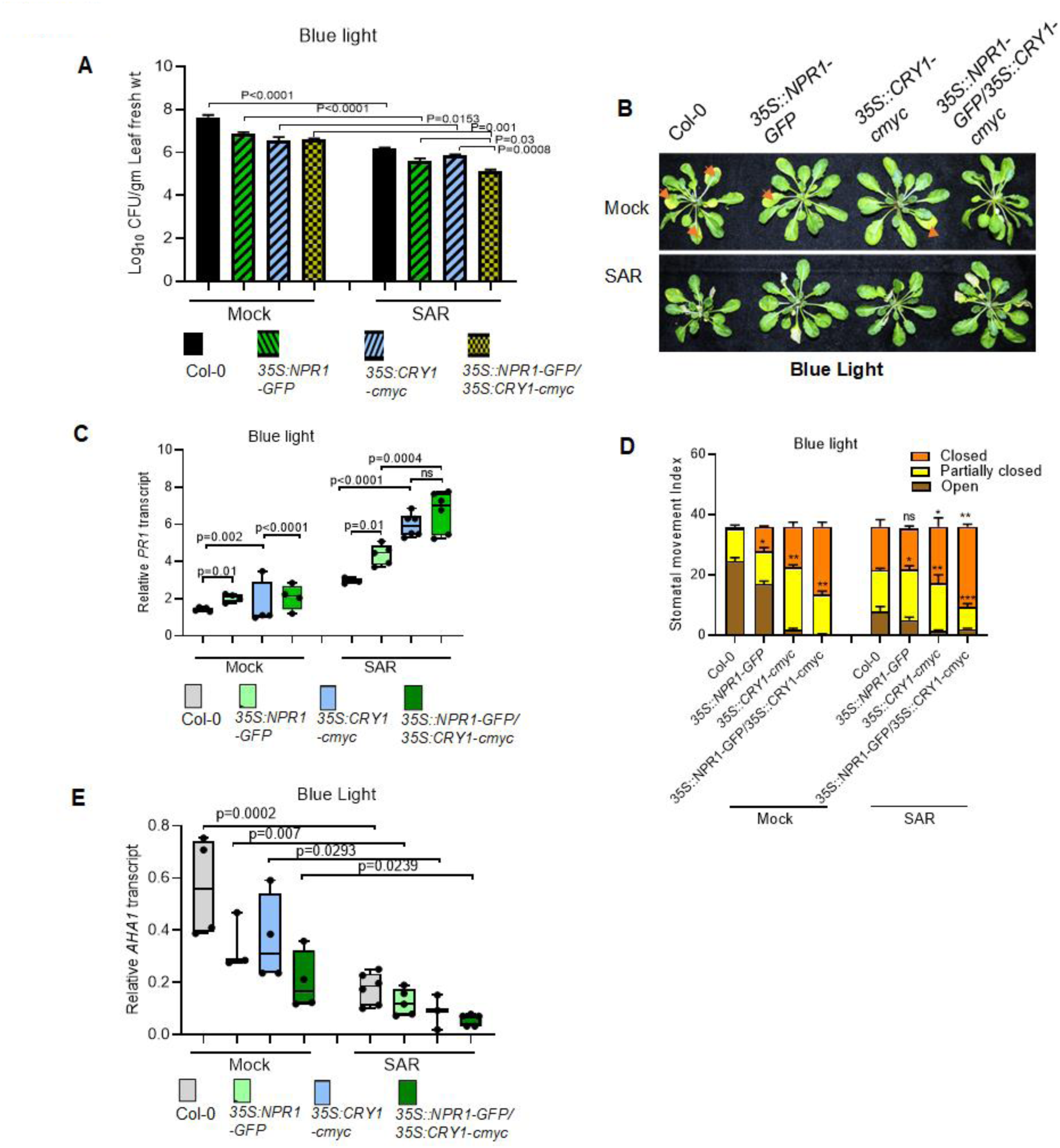
Plants overexpressing NPR1 and CRY showed SAR resistant phenotype under white and blue light associated with altered stomatal movement. (A) bacterial count assay in Col-0, *35S::NPR1-GFP*, *35S::CRY1-cmyc*, and *35SNPR1-GFP/35S-CRY1-cmyc* as described in the previous experiments under blue light. Bacterial load under white light is in the supplementary fig 5 A (B) disease symptom phenotype under blue light. Bacterial numbers were determined at 3 days post-infiltration (dpi). Each bar indicates the mean standard deviation (SD) (n = 4), as p values above the compared genotype, as obtained by two-way ANOVA (Tukey’s multiple comparison test) (C) Relative PR1 transcript under blue light in the systemic tissue of the shown genotypes. (D) stomatal movement index in the indicated genotypes. The stomatal movement index has been measured considering the typical phenotype of closed, Partially closed and opened stomata category as shown in the Figure 3. The stomatal movement index was measured under blue and white light conditions. Forty-eight hours after primary inoculation with mock or Avr, systemic leaves were detached and incubated in a stomatal opening solution for 2 hours to ensure full stomatal opening. The leaves were then transferred to a 5 µM flg22 solution. After 3 hours of flg22 treatment, stomata were examined using a confocal microscope. A total of 50 stomata per sample were counted and classified as open, partially open, or closed (as shown in G) across three independent biological replicates. (E) Relative *AHA1* expression in the distal tissue of secondary challenged leaves. Each bar indicates the mean standard deviation (SD) (n = 3) the significance has been denoted as p values above the compared genotype, as obtained by one-way ANOVA. Each bar indicates the mean standard deviation (SD) (n = 3 independent biological experiment) the significance has been denoted as p values above the compared genotype, as obtained by two-way ANOVA (Tukey’s multiple comparison test). The experiments repeated twice with similar results.

### SNRK3.6 phosphorylates CRY1 and is required for SAR under blue light

SnRK2.8 is known to be required for the phosphorylation and nuclear import of NPR1, which is associated with the activation of SAR (Lee et al., 2015b). We investigated for a similar mechanism for CRY1. A literature survey identified the nutrient-sensitive adenosine monophosphate–activated protein kinase (AMPK) that phosphorylates and promotes the degradation of cryptochrome 1 (CRY1) in mammalian cells and allows cryptochrome to relay nutrient signals to circadian clocks in mammalian peripheral organs like brain (Lamia et al., 2009; Lee and Kim, 2013). The ortholog of AMPK in Arabidopsis is SnRK3.6 (SNF1-Related Protein Kinase 3.6).

We first analyzed the expression profile of *SnRK3.6* under blue light treatment and assessed its transcript abundance in systemic tissue of wild type (Col-0) under blue light after 24 hours from a secondary challenge. Notably, *SnRK3.6* expression was found dependent on blue light over a 24-hour period (Figure 6A) and is also induced in systemic tissue during SAR development under blue light (Figure 6C) but downregulated under white light (Figure 6B). To explore the role of SnRK3.6 in developing SAR, the homozygous T-DNA insertion *snrk3.6* mutants (Supplemenatry Figure 5C) were exposed to a secondary infection by *Pst* DC3000 in systemic tissues after a primary infection with AvrRpt2 under blue-light conditions and bacterial count was measured. The *snrk3.6* mutants showed no restriction of bacterial growth and failed to develop systemic acquired resistance (SAR) under this condition (Figure 6D). To investigate any potential interaction between SnRK3.6 and CRY1, we transiently co-expressed them in *N. benthamiana* leaves for co-immunoprecipitation (Co-IP) (Figure 6E) and bimolecular fluorescence complementation (BiFC) assays (Figure 6F). Both the assays confirmed the *in vivo* interaction, with BiFC results further revealing that SnRK3.6 interacts with CRY1 at the plasma membrane and the nucleus (Figure 6F).

**Figure 6:**
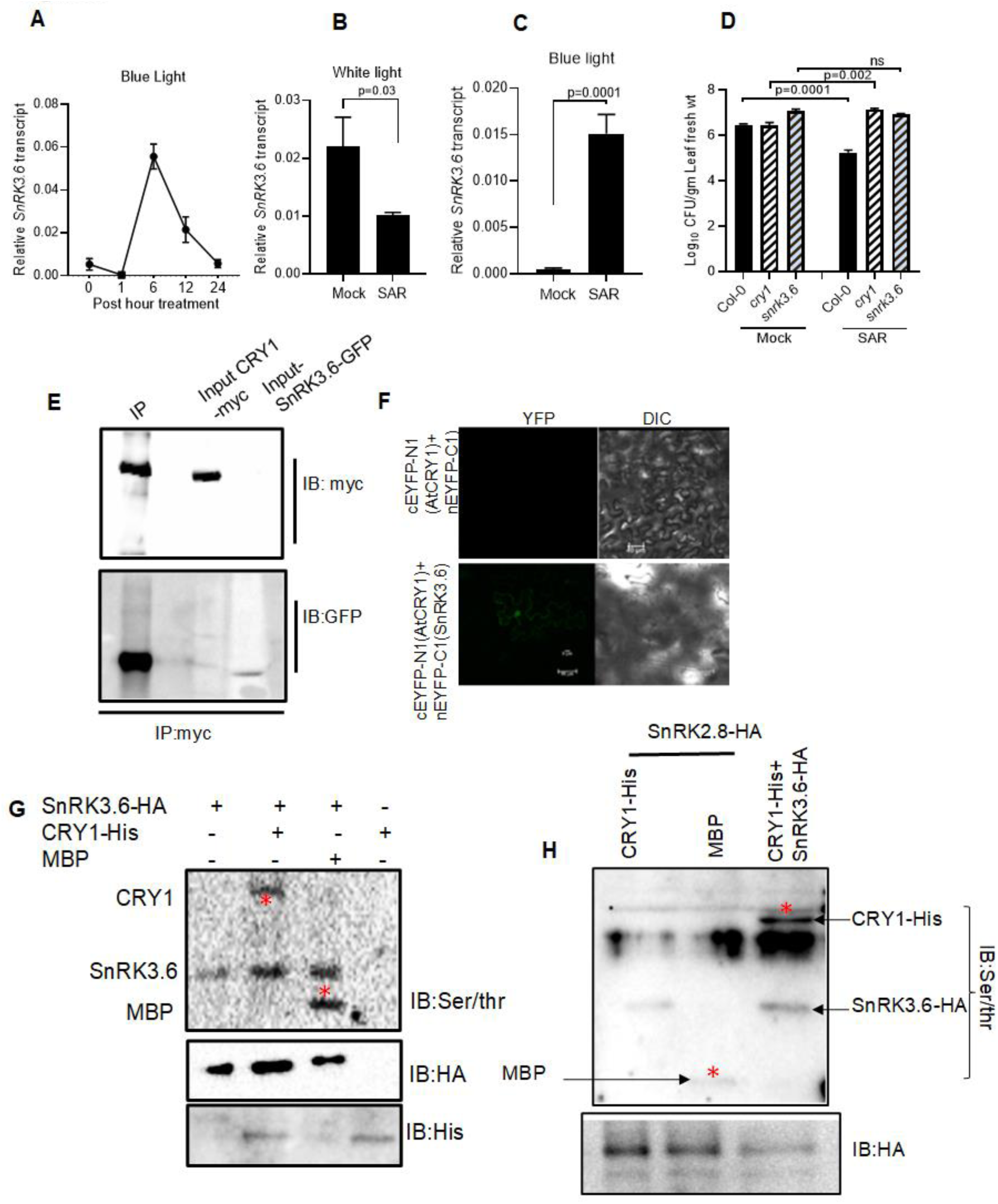
SNRK3.6 interacts with and phosphorylate CRY1 (A), (B), and (C) Transcript accumulation of SnRK3.6 under blue light and in mock and SAR tissue under white and blue light respectively. Bacterial numbers were determined at 3 days post-infiltration (dpi). Each bar indicates the mean standard deviation (SD) (n = 4, independent biological samples), as p values above the compared genotype, as obtained by two-way ANOVA (Tukey’s multiple comparison test). Mock (primary treatment =10 mM MgCl2 and secondary treatment = *Pst*DC3000); SAR (primary treatment = Avr-*Pst* and secondary treatment = *Pst*DC3000). (D) Bacterial count in the systemic tissue of the indicated genotypes under blue light. Each bar indicates the mean standard deviation (SD) (n = 4), as p values above the compared genotype, as obtained by two-way ANOVA (Tukey’s multiple comparison test) (C) Relative PR1 transcript under blue light in the systemic tissue of the shown genotypes. (E) Co-IP showing Interaction between CRY1 and SnRK3.6 *in planta.*(F) BIFC showing the interaction between CRY1 and SnRK3.6 (G*) In planta* phosphorylation of CRY1 through immunoprecipitated SnRK3.6-HA in Arabidopsis, red star shows the CRY1 or MBP detected through Ser/Thr antibody. (H) *In planta* phosphorylation of CRY1 through immunoprecipitated SnRK3.6-HA or SnRK2.8-HA in Arabidopsis, red star shows the CRY1 or MBP detected through Ser/Thr antibody. Immunoprecipitated SnRK2.8-HA did not phosphorylate CRY1-HIS.

SnRK3.6 is a serine/threonine protein kinase that encodes CBL-binding serine/threonine protein kinase 20 (CIPK20) featuring an N-terminal catalytic domain resembling SNF1/AMPK and a specific C-terminal regulatory domain that binds CBL. It regulates microtubule stability to mediate stomatal closure (Lee and Kim, 2013; Li et al., 2024). Interaction between CRY1 and SnRK3.6 prompted us to investigate whether SnRK3.6 phosphorylates CRY1. We performed an *in vitro* kinase assay by using bacterially purified recombinant CRY1 and SnRK3.6, and also by incubating bacterially purified CRY1 with immunoprecipitated SnRK3.6-HA from transiently expressing *N. benthamiana*. Phosphorylation was detected using a p-serine/p-threonine antibody, confirming SnRK3.6 phosphorylates CRY1 (Figure 6G, Supplementary Fig. 5A) while, SnRK2.8 and SnRK1.1 were unable to phosphorylate CRY1 (Figure 6H and Supplementary Fig. 5B) indicating that phosphorylation of CRY1 by SnRK3.6 was specific.

To understand the importance of SnRK3.6 and CRY1 interaction in the subcellular distribution of CRY1, we analyzed distribution of CRY1-GFP in plasma membrane, cytoplasmic, and nuclear fractions in CRY1-GFP-overexpressing wild-type (Col-0) and *snrk3.6* mutant plants. CRY1 was fractionated predominantly in the cytoplasm and nucleus in the wild-type background. In contrast, CRY1 accumulated mainly at the plasma membrane in the *snrk3.6* mutant. Nuclear localization of CRY1 was significantly higher in CRY1-GFP-overexpressing Col-0 than in the *snrk3.6* mutant (Figure 7A). Predominantly plasma membrane localization of CRY1-GFP in the *snrk3.6* background as compared to its wt counterpart was also visualized in the protoplasts isolated from these plants (Figure 7B). Our results described above suggests phosphorylation of CRY1 by SnRK3.6 is vital for its cellular distribution between plasma membrane, cytoplasmic and nuclear fractions. CRY1 localizes predominantly to the plasma membrane in absence of functional SnRK3.6 indicating that SnRK3.6 plays a key regulatory role in CRY1-mediated SAR response.

**Figure 7:**
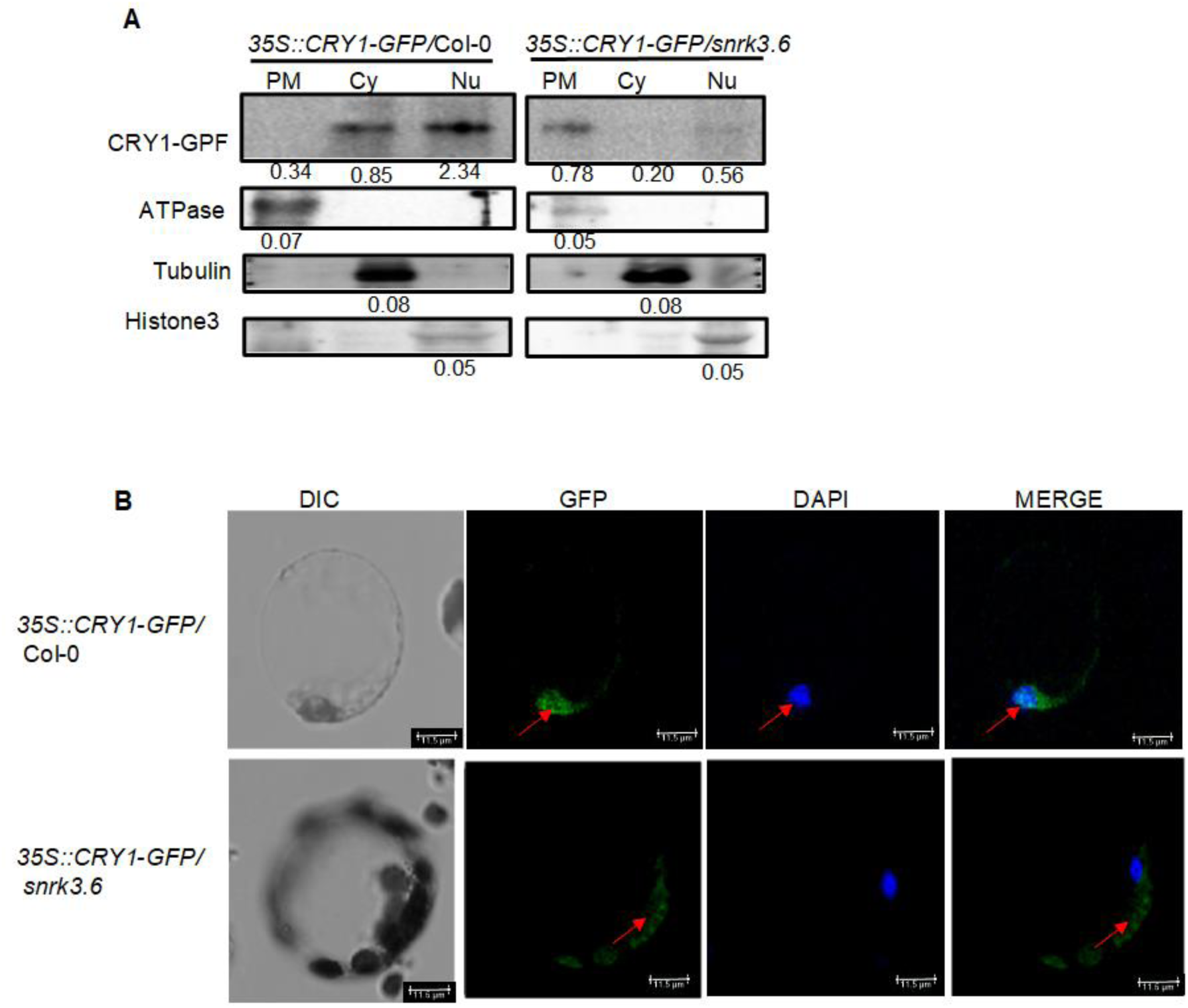
CRY1 localizes more to the nucleus in the presence of SnRK3.6. (A) A subcellular fractionation assay was performed in *Arabidopsis*, comparing two genetic backgrounds: Col-0 expressing CRY1-GFP and the *snrk3.6* mutant expressing CRY1-GFP. Equal amounts of protein (500 µg) from the plasma membrane, cytoplasmic, and nuclear fractions were used for immunoprecipitation (IP) with a GFP antibody. Following IP, Western blot analysis was carried out using the GFP antibody to detect and compare the levels of CRY1-GFP in each of the three subcellular compartments. To confirm equal protein loading and the purity of each fraction, ATPase (a plasma membrane marker), tubulin (a cytoplasmic marker), and Histone H3 (a nuclear marker) antibodies were used on the respective fractions.(B) To visualize and confirm the subcellular localization in the presence of SnRK3.6, protoplasts from the plants expressing CRY1 GFP in Col-0 and snrk3.6 mutant background were isolated and seen under confocal microscope

## Discussion

Plants adopt various immunomodulatory mechanisms to combat repeated pathogen attacks in the backdrop of continuously changing environment (Jian et al., 2024). Microenvironment play an integral role in this mechanistic regulation. Apart from its role in photosynthesis and development, light plays a major role in immune response in plants. This has been demonstrated using the photoreceptor mutants grown under varying light intensities nad wavelength (Kangasjärvi et al., 2012; Mawphlang and Kharshiing, 2017; Paik and Huq, 2019). Molecules involved in light perception, development and immune response function in an interconnected manner across the different biological system. It is well established that light activated CRY1 physically interacts with COP1 (CONSTITUTIVE PHOTOMORPHOGENIC 1) and SPA (SUPPRESSOR OF PHYTOCHROME A) proteins to form a complex and regulate plant development through HY5 and PIFs under light (Saijo et al., 2003; Lian et al., 2011; Kim et al., 2017; Podolec and Ulm, 2018). CRY1 also play a major role in development through binding PIF4/5 and BZR1 and further repressing the auxin and brassinosteroid signalling under blue light (Oh et al., 2012; He et al., 2019). In a recent study, NPR1 has been shown to play in blue light dependent photomorphogenesis by facilitating degradation of PIF4 (Zhou et al., 2024). It reveals that NPR1 functions as molecular bridge not just in systemic acquired resistance (SAR) but also in light dependent photomorphogenesis.

In our study we tried to fill a critical knowledge gap between light perception and immune priming. Our study unambiguously demonstrated a direct molecular interaction between these two major pathways. Our results suggest that CRY1 functions as a light-dependent interactor of NPR1, ensuring effective systemic immunity under blue-light conditions. This coupling not only enriches our understanding of how plants induce SAR under shade or blue light, but also precisely illustrates intertwined control of growth, development, and defence under different light regimes in nature.

### CRY1 is essential for SAR under low-intensity/blue light

We found that *cry1* mutants failed to develop SAR in low intensity or blue light, supported by reduced *PR1*, *PR2*, *NPR1* expression in distal leaves resulting into a higher susceptibility to secondary *Pst*DC3000 infection. Under normal white light, *cry1* and wild type (Col-0) both mount SAR effectively. While CRY1 is classically linked to photomorphogenesis, many defence genes (like *PR1*, *NPR1*) are also regulated by light. Blue-light perception *via* CRY1 may prime or boost immune signaling, perhaps by modulating H₂O₂ levels, phytohormones, or defence genes. This complements our observations that *cry1* mutants poorly express PR genes under blue or low light. Further, at 26 °C under the same light condition, *cry1* mutants exhibited reduced bacterial load post-secondary challenge and robust *PR1*/*NPR1* induction, suggesting SAR is active. However, *cry1* mutants failed in inducing SAR and defence genes at 22 °C under same low light condition, indicating temperature critically modulates the SAR phenotype. Together, temperature plays a deciding role in the functioning of CRY1 in mounting SAR.

Apart from regulating plant defense through controlling stomatal aperture (Hao et al., 2025a), there could be other ways through which CRY1 can impact SAR. For example, down stream to CRY1, PIF4 mediates thermosensory growth of plants that results in disease resistance in Arabidopsis (Gangappa et al., 2017). The regulator of photomorphogenesis HY5 is also shown to be involved in defense induction NPR1 interacts with HY5 in a light-dependent way, which is disrupted by the bacterial effector protein AvrPtoB (Liu et al., 2025a). On the other hand, CRY1 also interacts with HY5 through COP1 to mediate blue light-mediated thermotolerance in which CRY1-mediated signaling suppresses thermotolerance (Liu et al., 2025b). CRY1 affects auxin, gibberellin, and possibly salicylic acid (SA) pathways these are central to SAR. CRY1 helps regulate circadian rhythms, which in turn gate immune responses temporally; disruption under low light could impair timely SAR. All the information mentioned above suggest that light and temperature-mediated regulation of plant defense is governed by overlapping pathways. The functional hypothesis connecting our results about the role of CRY1 in low intensity/blue light could be that CRY1 after getting phosphorylated by SnRK3.6 assists NPR1 to move to nucleus and/or CRY1 likely shifts photochemically into a signaling-competent state that primes SAR via SA/NPR1/PR pathways. In *cry1* mutants, absence of this active state may unlock alternative defense regulators, allowing SAR induction through compensatory pathways (e.g. other photoreceptors, circadian modulators) at higher growth temperature at low light. Whereas at normal growth temperature under low light, CRY1 is less active or required to maintain transcriptional activation (via HY5, PIF4) and, therefore, *cry1* mutants cannot mount SAR effectively.

### NPR1 is partially dispensable, but full SAR requires CRY1–NPR1 interaction and NPR1-CRY1 interaction is light dependent and nuclear

Our data show that blue light enhances both NPR1 transcript and protein levels, which aligns with the well-established role of blue-light photoreceptors in modulating nuclear-localized transcription factors *via* COP1/HY5 and related pathways. This suggests that CRY1 is upstream of NPR1 in a blue-light signaling cascade, enabling more robust SAR induction under these conditions. The observation that *npr1-2* mutants can mount SAR under blue light despite their known SAR deficits under full-spectrum, high-intensity light implies two possibilities: First, there is alternate blue-light–driven immune pathways that may partially compensate the absence of NPR1, perhaps involving photoreceptor-driven stomatal defense or ROS-mediated mechanisms. Second, stomatal control becomes a dominant factor when blue light is the primary stimulus, consistent with studies showing that CRY1 can modulate stomatal innate immunity *via* interactions with FLS2 and components like LURP1 (Hao et al., 2025b).

Our stomatal index measurement experiment revealed that under white light, *cry1* and wild type (Col-0) show similar stomatal response patterns, but *npr1-2* has increased partially closed stomata consistent with known role of NPR1 in guard-cell signaling. Whereas under blue light, *cry1* had increased open and partially closed stomata as compared to wild type (Col-0), suggesting impaired stomatal closure due to lack of CRY1-mediated signaling. The *npr1-2* mutants showed greater number of closed stomata indicating towards an enhance stomatal defense phenotype during SAR induction. This supports the notion that blue light potentiates stomatal closure in *npr1* mutants, perhaps *via* CRY1-independent routes, thereby reducing pathogen colonization and enabling a form of SAR. We propose that in Col-0 under blue light, CRY1 stabilizes COP1/HY5 and other transcription factors, which in turn induce *NPR1* expression and activity, facilitating classical *PR* gene-mediated SAR. For *npr1-2* under blue light, alternatively regulated stomatal defense pathways e.g., increased ROS production or stomatal closure *via* LURP1(Late Up-regulation in response to Hpa1) /FLS2 (Flagellin sensitive 2) (Hao et al., 2025c) complexes might compensate to limit pathogen spread, even without PR gene induction. In *cry1* mutants, blue light fails to activate stomatal immunity and NPR1 induction, leading to open stomata and compromised SAR. Thus, blue light signaling may bifurcates into transcriptional SAR *via* CRY1→NPR1→PR genes; and stomatal immunity likely *via* CRY1-independent but blue light–responsive modules (e.g., LURP1/FLS2). Loss of CRY1 disables both, but loss of NPR1 disables only the transcriptional arm leaving stomatal closure intact and permitting SAR under blue light.

To dissect the molecular interplay between NPR1 and CRY1 in blue light–mediated systemic acquired resistance (SAR), we employed a suite of biochemical and cell-based assays complemented by *in planta* validation. An *in vitro* GST pull-down confirmed a direct physical interaction between NPR1 and CRY1. This association was corroborated *in vivo* through split-luciferase complementation in *Nicotiana benthamiana*, where NPR1-nLUC and CRY1-cLUC reconstituted luciferase activity. Bimolecular fluorescence complementation localized this interaction predominantly to the nucleus. Building on this, co-immunoprecipitation (Co-IP) followed by immunoblot showed a marked enhancement of the NPR1-CRY1 interaction under blue light in comparison to that under white light. Fractionation followed by Co-IP further revealed higher nuclear abundance of NPR1 and CRY1 in systemic tissues under blue light, with NPR1 accumulating more than CRY1. These data suggest that blue light promotes nuclear assembly of the NPR1-CRY1 complex to regulate SAR. This aligns with prior studies demonstrating that cryptochromes, upon blue light activation, undergo conformational changes to enable nuclear interactions with signaling partners. Together, our results highlight a blue light–dependent, nuclear-localized NPR1-CRY1 complex as a pivotal node in linking photoreceptor activity to systemic immune signaling. Also, The enhanced nuclear accumulation of both NPR1 and CRY1 under blue light suggests a synergistic interaction between these proteins in mounting SAR. The preferential nuclear localization of NPR1 over CRY1 may indicate a primary role for NPR1 in initiating the transcriptional responses required for SAR, with CRY1 potentially modulating these responses through its light-dependent activities. This study advances our understanding of the molecular mechanisms underlying SAR and highlights the complex interplay between light signaling and immune responses in plants.

Our data also demonstrate that overexpression of CRY1 alone improves systemic acquired resistance (SAR) under blue light, reducing bacterial load compared to Col-0. When combined with NPR1 overexpression, this effect is amplified, yielding a stronger SAR phenotype and higher *PR1* transcript levels in systemic tissues. This aligns with the broader role of CRY1 in enhancing SA- and light-mediated defense responses particularly blue-light-induced SAR, where CRY1 has been recognized as the principal photoreceptor mediating this response. *NPR1* overexpression triggers *PR1* induction in both local and systemic tissues across as expected given central role of NPR1 as the SA-receptor and transcriptional co-activator of *PR* genes. Whereas, *CRY1* overexpression under blue light induces NPR1 expression in systemic tissues which likely elevates NPR1 abundance or activation highlighting a positive feedback where CRY1 amplifies the SA and NPR1 mediated defense.

Stomata are key openings for bacterial pathogens (Melotto et al., 2008). NPR1 alone increases partial stomatal closure; CRY1 further induces both partial and full closure. The double overexpressor shows the highest full closure rate. This phenotype correlates with significantly reduced transcript levels of AHA1 (H⁺-ATPase), which is essential for stomatal opening. This is consistent with a mechanism where CRY1 enhances NPR1-driven stomatal defense under blue-light conditions, further limiting bacterial entry.

### SnRk3.6 is a new connector

In mammals, AMPK phosphorylates CRY1, promoting its degradation and linking nutrient status to circadian rhythms. The Arabidopsis ortholog SnRK3.6 (OST1) is thus a prime kinase candidate for phosphorylating CRY1, especially since AMPK-related SnRKs are known to mediate stress and hormone signaling. Our results that SnRK3.6 expression is enhanced by blue light and during SAR highlights a light-responsive regulatory mechanism. This expression profile indicated a possible role for SnRK3.6 to act as a blue-light-activated kinase that may modulate CRY1 activity and/or localization during immune responses. The *snrk3.6* mutants are insensitive to secondary challenge to induce SAR. Co-IP and BiFC data demonstrate SnRK3.6 physically interacts with CRY1 at the plasma membrane. SnRK3.6 (also known as CBL-interacting protein kinase 20, AT5G45820) is known to bind CBL proteins. SnRK3 familiy proteins were previously shown to regulate ion homeostasis and stomatal closure (Mondal et al., 2024). A membrane-associated SnRK3.6-CRY1 complex suggests signaling initiation at the cell periphery, potentially linking guard-cell dynamics and SA transport to CRY1-mediated SAR. *In vitro* kinase assays confirmed that CRY1 is specifically phosphorylated by SnRK3.6. This mirrors the mammalian model where AMPK specifically phosphorylates CRY1 (Lamia et al., 2009). SnRK3.6 interacts with CRY1 at the plasma membrane and likely influences the nuclear localization of CRY1 by modulating its post-translational state, stability, and/or signaling function under blue light. Our subcellular fractionation assay revealed that a substantial portion of CRY1 shifts to the nucleus. In *snrk3.6* mutants, nuclear accumulation of CRY1 is significantly reduced, with more protein sequestered in membrane or cytosolic fractions This supports the idea that SnRK3.6 is required for maximum nuclear translocation of CRY1, possibly via its phosphorylation.

In conclusion, based on our experimental evidence, we proposed a model on SAR operates efficiently under blue light or low-light conditions and how CRY1 and its modulating kinase play a major role in that. Blue light serves as a key signal that activates the photoreceptor CRY1, along with the kinases SnRK3.6. Previous studies reported the role of SnRK2.8 in phosphorylating and monomerizing NPR1, which facilitates its nuclear transport in the systemic tissues where it initiates defense gene expression. Simultaneously, CRY1 is also activated and may be phosphorylated in response to blue light through SnRK3.6. Once activated, CRY1 physically interacts with NPR1 in systemic tissues following a secondary infection. This CRY1–NPR1 complex may enhance the expression of pathogenesis-related (*PR*) genes and *NPR1* itself, while also promoting stomatal defense by inducing *AHA1* expression and facilitating stomatal closure. Collectively, these processes strengthen stomatal immunity, limiting pathogen entry and improving overall plant defense responses under blue light conditions (Figure 8).

**Figure 8:**
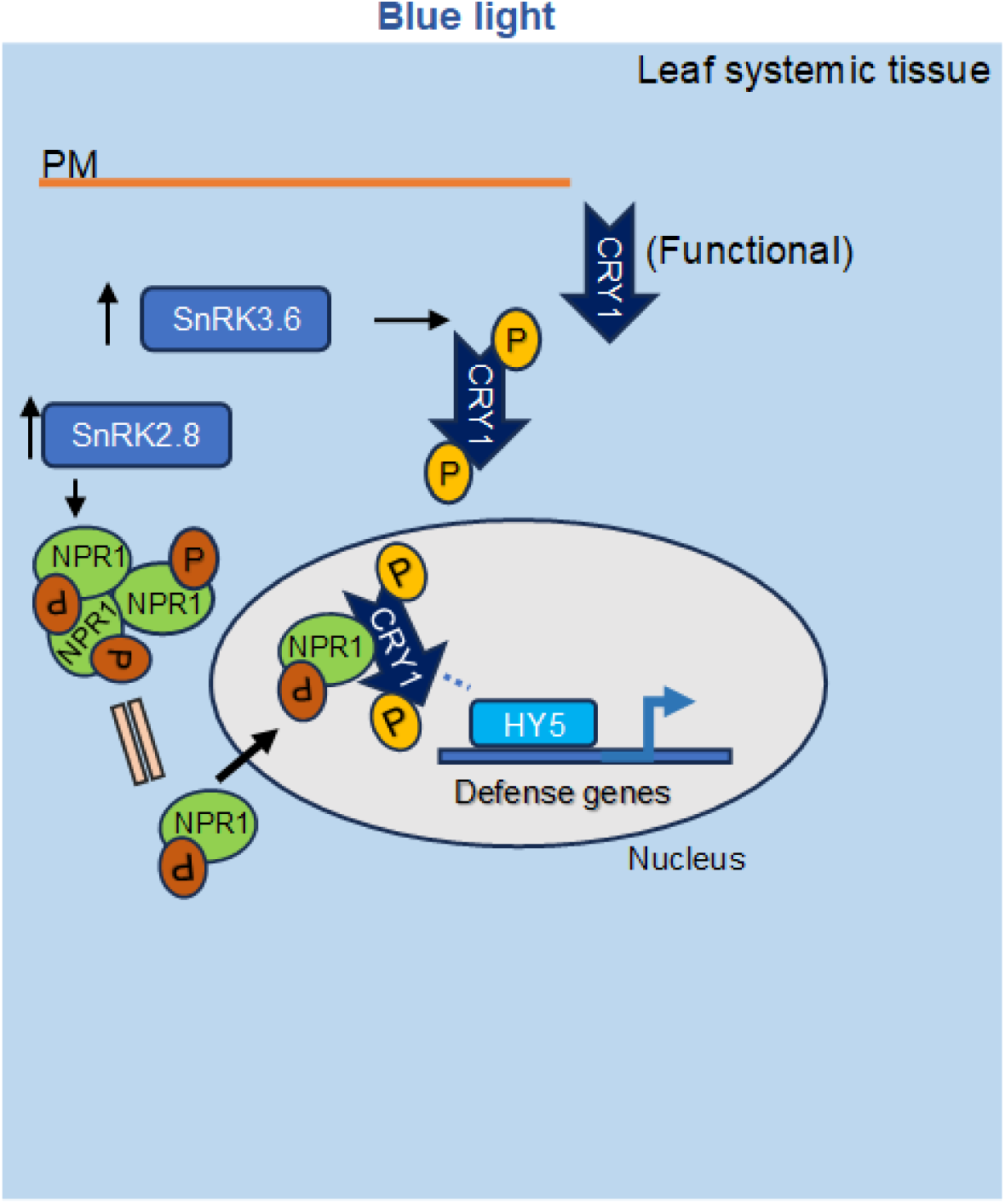
Proposed Model: Enhanced SAR (Systemic Acquired Resistance) is effective in blue/light-limited conditions and not functional in *cry1* mutants under blue light. Blue light activates the CRY1, SnRK3.6, NPR1, and SnRK2.8. SnRk2.8 phosphorylates NPR1, converts its polymeric form to monomeric form that in turn localises into the nucleus in the systemic tissue. Similarly, CRY1 is activated in blue light and phosphorylated by SnRK3.6. The CRY1 physically interacts with NPR1 under blue light in the systemic tissue after *Pst*DC3000 challenge. CRY1-NPR1 complex promotes *PR* genes and *NPR1* expression, stomatal defense via synergistic closure and *AHA1* expression it results in improved stomatal immunity and prevents pathogen entry.

## Materials and Methods

### Plant material and growth conditions

All of the Arabidopsis genotypes were in Col-0 ecotype. The mutants *npr1-2*, *phyA*, *phyB* and *cry1*, and Col-0 have been used from our lab stocks. The overexpression plants of NPR1(*35S::NPR1-GFP*/*myc*) and CRY1(*35S::CRY1*-*GFP*/*myc*) were generated in their mutant backgrounds. The homozygous plants were confirmed through RT-PCR and western blot with either GFP or myc antibody. The homozygous CRY1-myc and NPR1-GFP were utilised for crossing to develop homozygous CRY1-myc/NPR1-GFP plants and they were confirmed through western blot. 35S::CRY1-GFP/*snRk3.6* plants were generated through agrobacterium mediated floral dip transformation of *snrk3.6* (Salk_040637C, obtained from ABRC) homozygous mutants, confirmed through PCR, using Agrobacterium harboring pGWB6-35SCRY1-GFP. The T_3_ homozygous lines were confirmed through RT-PCR for checking the expression of CRY1.The plants were grown in controlled growth condition at 22^0^C, 60% relative humidity, and photoperiod of 8hrs light and 16 hrs dark cycle.

### *Pseudomonas syringae* infection assay for systemic resistance

SAR assays were conducted following previously established protocols (Singh et al., 2013; Singh et al., 2023). Briefly, primary inoculation was done with over-night grown *Avr-Pst* culture. The Avr-Pst was infiltrated into the lowermost two to three leaves of 5-week-old plants at a concentration of 10⁷ CFU/ml. Plants infiltrated with 10 mM MgCl₂ served as mock controls. After three days of primary inoculation, distal/ upper leaves were challenged with *Pseudomonas syringae* pv. *tomato* (*Pst*DC3000) at 10⁵ CFU/ml. Bacterial counts were assessed three days secondary post-inoculation (dpi) by plating serial dilutions of homogenized leaf tissue on rifampicin and kanamycin containing TSGA (tryptone-sucrose-Na-L glutamate-Agar) media. For each plant genotype, four biological replicates were analyzed, with each replicate consisting of five leaf collected from different plants.

### Stomatal aperture measurement

Epidermal layer peeled off from the abaxial side of the Arabidopsis leaves for all the genotypes kept under white and blue light, floated in the stomatal opening buffer for 2-2.5hrs. The primary leaves were inoculated and the distal leaves tissue were used for stomatal aperture measurement. The distal leaves treated with *flg22* for 2-3 hrs were incubated in a stomatal opening solution for 2–3 hours and observed under a light microscope. 50-75 stomata were randomly selected for measuring the aperture using ImageJ software (Gupta et al., 2020; Gupta and Nandi, 2020).

### RNA extraction and qPCR analysis

Total RNA from Arabidopsis leaves were isolated by Trizol (Sigma) method. DNase treatment was done to remove DNA before making cDNA. cDNA was made from 1 µg of RNA by Thermo Verso cDNA synthesis kit. The qPCR was done by using 50-100 ng of cDNA and power SYBR green master mix (Applied Biosystem, USA) in ABI-PRISM 7500 FAST sequence detector. For qPCR analysis, typically each sample contained three biological replicates. qPCR was carried out by taking two technical replicates of each biological cDNA (Singh and Nandi, 2022). The primers used in the expression analysis are enlisted in supplementary table S1.

### In vitro pull-down assay

The NPR1 and CRY1 coding sequence was amplified and cloned in pGEX4T2 and pET28a^+^ for recombinant protein expression. The respective proteins were expressed in BL21-codon^+^ E.coli cells. The GST-NPR1 fusion protein was purified through GST-sepharose beads via 50mM reduced glutathione whereas CRY1-His was purified through Ni-NTA beads via 200mM Imidazole. The purified proteins were incubated at 4^0^C on the rotating shaker at very low speed, further GST beads were added and incubated overnight at 4^0^C on the rotating shaker. Beads were spun down at 150g, removed the supernatant and further washed with washing buffer containing 5mM reduced glutathione, finally the bound proteins were eluted through 50mM reduced glutathione via incubating at 4^0^C for 15 minutes. The eluted proteins were run on the SDS page and detected through western blot using GST and HIS antibody for NPR1-GST and CRY1-His respectively.

### Co-IP

Coimmunoprecipitation has been done as described in Singh et al., 2018 (Singh et al., 2018). in the Arabidopsis plants expressing both NPR1-GFP and CRY1-myc. The 4-week-old Arabidopsis plants kept under blue and white light were primary challenged with AvrRpt2 followed by *Pst*DC3000 inoculation in the distal tissue. After 48 hrs of secondary challenge distal leaves tissue have been harvested for total protein extraction. The total protein has been first incubated with myc antibody for 2hrs at 4^0^C on the rotating shaker at very low speed. 50ul of DYNAbeads^TM^ proteinG were added in the total protein and myc antibody mix and incubated for 4-6 hrs at 4^0^C on the rotating shaker. The supernatant has been discarded and beads have been washed 3-4 times with 1XPBS/TBS containing 0.1% tween-20. The washed beads have been run on 10% SDS-PAGE. The respective corresponding to NPR1 and CRY1 were detected through western blot developed through GFP and cmyc antibody.

### BIFC

For performing the BiFC assay, full length coding sequence of NPR1 was amplified and fused with N-terminal YFP fragment of CD3-1648 (pSITE-nEYFP-C1) plasmid vector to generate MNPR1-nEYFP fusions and CRY1 CDS was amplified and fused with C-terminal YFP fragment in CD3-1651 (pSITE-cEYFP-N1) plasmid vector to make CRY1-cEFP fusion construct. The constructed plasmids were transformed in Agrobacterium tumefaciens and transiently expressed in *Nicotiana benthamiana*. The fluorescence was observed after 48h of infiltration under confocal microscope at the same focus and exposure of laser.

### Split luciferase assay

To check the interaction of NPR1 And CRY1 in Nicotiana benthamiana the NPR1 was clone in pCAMBIA1300-nLuc to make the nluc-NPR1 fusion and CRY1 was cloned in pCAMBIA1300-cLuc to make the CRY1-cLuc fusion. Both the constructs were transiently co-expressed in *N.benthamiana* leaves, the single construct (pCAMBIA1300-nLuc-NPR1) and the only vector (pCAMBIA1300-cLuc) were utilised as negative controls. The higher the intensity of luminescence in the pCAMBIA1300-nLuc- NPR1 and pCAMBIA1300-CRY1-cLuc co-expressing leaves as compared to pCAMBIA1300-nLuc and pCAMBIA1300-CRY1-cLuc shows the positive interaction between NPR1 and CRY1 *in* planta.

## Acknowledgements and fundings

The authors acknowledge the funding resources provided by the BRIC-National Institute of Plant Genome Research, Department of Biotechnology (DBT), Ministry of Science and Technology, Government of India. DC acknowledges J.C. Bose Fellowship (JCB/2020/000014) from Science and Engineering Research Board, Department of Science and Technology. NS was supported by CSIR-SRA-pool scientist scheme, Award No. 13(9170-A)/2021-Pool by Council of Scientific and Industrial Research, Govt. of India.

## Authors contributions

NS, DC, and SC initiated, conceived, designed, and coordinated the research project. SC and DC have given the intellectual inputs during the preparation of the MS. NS wrote the manuscript, SC and DC edited it. NS developed plant lines, reagents, performed experiments. GJ contributed in developing transgenic plant lines. NS, SC, and DC analyzed the data and finalized the manuscript.

Inducers of Plant Systemic Acquired Resistance Regulate NPR1 Function through Redox Changes.

## References

1. Abramovitch RB, Anderson JC, Martin GB (2006) Bacterial elicitation and evasion of plant innate immunity. Nat Rev Mol Cell Biol 7: 601–611

2. Backer R, Naidoo S, van den Berg N (2019a) The NONEXPRESSOR OF PATHOGENESIS-RELATED GENES 1 (NPR1) and related family: Mechanistic insights in plant disease resistance. Front Plant Sci. doi: 10.3389/fpls.2019.00102

3. Backer R, Naidoo S, van den Berg N (2019b) The NONEXPRESSOR OF PATHOGENESIS-RELATED GENES 1 (NPR1) and related family: Mechanistic insights in plant disease resistance. Front Plant Sci. doi: 10.3389/fpls.2019.00102

4. Ballaré CL, Mazza CA, Austin AT, Pierik R (2012) Canopy light and plant health. Plant Physiol 160: 145–155

5. Ballaré CL, Pierik R (2017) The shade-avoidance syndrome: Multiple signals and ecological consequences. Plant Cell Environ 40: 2530–2543

6. Blanco F, Garretón V, Frey N, Dominguez C, Pérez-Acle T, Van Der Straeten D, Jordana X, Holuigue L (2005) Identification of NPR1-dependent and independent genes early induced by salicylic acid treatment in arabidopsis. Plant Mol Biol 59: 927–944

7. Breen S, McLellan H, Birch PRJ, Gilroy EM (2023) Tuning the Wavelength: Manipulation of Light Signaling to Control Plant Defense. Int J Mol Sci. doi: 10.3390/ijms24043803

8. Briggs WR (2014) Phototropism: Some history, some puzzles, and a look ahead. Plant Physiol 164: 13–23

9. Cao H, Glazebrook J, Clarke JD, Volko S, Dong X (1997) The Arabidopsis NPR1 gene that controls systemic acquired resistance encodes a novel protein containing ankyrin repeats. Cell 88: 57–63

10. Cashmore AR, Jarillo JA, Wu Y-J, Liu D (1988) Cryptochromes: Blue Light Receptors for Plants and Animals. CRC Press

11. Chandra-Shekara AC, Gupte M, Navarre D, Raina S, Raina R, Klessig D, Kachroo P (2006) Light-dependent hypersensitive response and resistance signaling against Turnip Crinkle Virus in Arabidopsis. Plant Journal 45: 320–334

12. Christie JM, Blackwood L, Petersen J, Sullivan S (2015) Plant flavoprotein photoreceptors. Plant Cell Physiol 56: 401–413

13. da Cunha L, McFall AJ, Mackey D (2006) Innate immunity in plants: a continuum of layered defenses. Microbes Infect 8: 1372–1381

14. Gallé Á, Czékus Z, Tóth L, Galgóczy L, Poór P (2021) Pest and disease management by red light. Plant Cell Environ 44: 3197–3210

15. Gangappa SN, Berriri S, Kumar SV (2017) PIF4 Coordinates Thermosensory Growth and Immunity in Arabidopsis. Current Biology 27: 243–249

16. Genoud T, Buchala AJ, Chua NH, Métraux JP (2002a) Phytochrome signalling modulates the SA-perceptive pathway in Arabidopsis. Plant Journal 31: 87–95

17. Genoud T, Buchala AJ, Chua NH, Métraux JP (2002b) Phytochrome signalling modulates the SA-perceptive pathway in Arabidopsis. Plant Journal 31: 87–95

18. Guan Q, David L, Moran R, Grela I, Ortega A, Scott P, Warnock L, Chen S (2023) Role of NPR1 in Systemic Acquired Stomatal Immunity. Plants. doi: 10.3390/plants12112137

19. Guo A, Reimers PJ, Leach JE (1993) Effect of light on incompatible interactions between Xanthomonas oryzae pv oryzae and rice. Physiol Mol Plant Pathol 42: 413–425

20. Gupta P, Nandi AK (2020) Long-chain base kinase1 promotes salicylic acid-mediated stomatal immunity in Arabidopsis thaliana. J Plant Biochem Biotechnol 29: 796–803

21. Gupta P, Roy S, Nandi AK (2020) MEDEA-interacting protein LONG-CHAIN BASE KINASE 1 promotes pattern-triggered immunity in Arabidopsis thaliana. Plant Mol Biol 103: 173–184

22. Hao Y, Zeng Z, Yuan M, Li H, Guo S, Yang Y, Jiang S, Hawara E, Li J, Zhang P, et al (2025a) The blue-light receptor CRY1 serves as a switch to balance photosynthesis and plant defense. Cell Host Microbe 33: 137–150.e6

23. Hao Y, Zeng Z, Yuan M, Li H, Guo S, Yang Y, Jiang S, Hawara E, Li J, Zhang P, et al (2025b) The blue-light receptor CRY1 serves as a switch to balance photosynthesis and plant defense. Cell Host Microbe 33: 137–150.e6

24. Hao Y, Zeng Z, Yuan M, Li H, Guo S, Yang Y, Jiang S, Hawara E, Li J, Zhang P, et al (2025c) The blue-light receptor CRY1 serves as a switch to balance photosynthesis and plant defense. Cell Host Microbe 33: 137–150.e6

25. He G, Liu J, Dong H, Sun J (2019) The Blue-Light Receptor CRY1 Interacts with BZR1 and BIN2 to Modulate the Phosphorylation and Nuclear Function of BZR1 in Repressing BR Signaling in Arabidopsis. Mol Plant 12: 689–703

26. Jeong RD, Chandra-Shekara AC, Barman SR, Navarre D, Klessig DF, Kachroo A, Kachroo P (2010) Cryptochrome 2 and phototropin 2 regulate resistance protein-mediated viral defense by negatively regulating an E3 ubiquitin ligase. Proc Natl Acad Sci U S A 107: 13538–13543

27. Jian Y, Gong D, Wang Z, Liu L, He J, Han X, Tsuda K (2024) How plants manage pathogen infection. EMBO Rep 25: 31–44

28. Jones JDG, Dangl JL (2006) The plant immune system. Nature 444: 323–329

29. Kangasjärvi S, Neukermans J, Li S, Aro EM, Noctor G (2012) Photosynthesis, photorespiration, and light signalling in defence responses. J Exp Bot 63: 1619–1636

30. Katagiri F, Thilmony R, He SY (2002) The Arabidopsis Thaliana-Pseudomonas Syringae Interaction. Arabidopsis Book 1: e0039

31. Kaur S, Samota MK, Choudhary M, Choudhary M, Pandey AK, Sharma A, Thakur J (2022) How do plants defend themselves against pathogens-Biochemical mechanisms and genetic interventions. Physiology and Molecular Biology of Plants 28: 485–504

32. Kim JY, Song JT, Seo HS (2017) COP1 regulates plant growth and development in response to light at the post-translational level. J Exp Bot 68: 4737–4748

33. Klessig DF, Choi HW, Dempsey DA (2018) Systemic acquired resistance and salicylic acid: Past, present, and future. Molecular Plant-Microbe Interactions 31: 871–888

34. Lamia KA, Sachdeva UM, Di Tacchio L, Williams EC, Alvarez JG, Egan DF, Vasquez DS, Juguilon H, Panda S, Shaw RJ, et al (2009) AMPK regulates the circadian clock by cryptochrome phosphorylation and degradation. Science (1979) 326: 437–440

35. Lee HJ, Park YJ, Seo PJ, Kim JH, Sim HJ, Kim SG, Park CM (2015a) Systemic immunity requires SnRK2.8-mediated nuclear import of NPR1 in arabidopsis. Plant Cell 27: 3425–3438

36. Lee HJ, Park YJ, Seo PJ, Kim JH, Sim HJ, Kim SG, Park CM (2015b) Systemic immunity requires SnRK2.8-mediated nuclear import of NPR1 in arabidopsis. Plant Cell 27: 3425–3438

37. Lee Y, Kim EK (2013) AMP-activated protein kinase as a key molecular link between metabolism and clockwork. Exp Mol Med. doi: 10.1038/emm.2013.65

38. Li T, Zhou X, Wang Y, Liu X, Fan Y, Li R, Zhang H, Xu Y (2024) AtCIPK20 regulates microtubule stability to mediate stomatal closure under drought stress in Arabidopsis. Plant Cell Environ. doi: 10.1111/pce.15112

39. Lian HL, He SB, Zhang YC, Zhu DM, Zhang JY, Jia KP, Sun SX, Li L, Yang HQ (2011) Blue-light-dependent interaction of cryptochrome 1 with SPA1 defines a dynamic signaling mechanism. Genes Dev 25: 1023–1028

40. Liu J, van Iersel MW (2021) Photosynthetic Physiology of Blue, Green, and Red Light: Light Intensity Effects and Underlying Mechanisms. Front Plant Sci. doi: 10.3389/fpls.2021.619987

41. Liu P, Zhao Z, Tang Y, Zhou Y, Liu J, Xu K, Chen Y, Li X, Tang Y, Yang L (2025a) The HY5-NPR1 module governs light-dependent virulence of a plant bacterial pathogen. Cell Host Microbe 33: 1606–1622.e10

42. Liu S, Wang Q, Zhong M, Lin G, Ye M, Wang Y, Zhang J, Wang Q (2025b) The CRY1–COP1–HY5 axis mediates blue-light regulation of Arabidopsis thermotolerance. Plant Commun. doi: 10.1016/j.xplc.2025.101264

43. Malinovsky FG, Fangel JU, Willats WGT (2014) The role of the cell wall in plant immunity. Front Plant Sci. doi: 10.3389/fpls.2014.00178

44. Mawphlang OIL, Kharshiing E V. (2017) Photoreceptor mediated plant growth responses: Implications for photoreceptor engineering toward improved performance in crops. Front Plant Sci. doi: 10.3389/fpls.2017.01181

45. Melotto M, Underwood W, Sheng YH (2008) Role of stomata in plant innate immunity and foliar bacterial diseases. Annu Rev Phytopathol 46: 101–122

46. Mondal S, Paul A, Mitra D, Pradhan C, Seth CS, Chattopadhyay K, Chakraborty K (2024) The multifaceted role of different SnRK gene family members in regulating multiple abiotic stresses in plants. Physiol Plant. doi: 10.1111/ppl.14543

47. Naqvi S, He Q, Trusch F, Qiu H, Pham J, Sun Q, Christie JM, Gilroy EM, Birch PRJ (2022) Blue-light receptor phototropin 1 suppresses immunity to promote Phytophthora infestans infection. New Phytologist 233: 2282–2293

48. Oh E, Zhu JY, Wang ZY (2012) Interaction between BZR1 and PIF4 integrates brassinosteroid and environmental responses. Nat Cell Biol 14: 802–809

49. Paik I, Huq E (2019) Plant photoreceptors: Multi-functional sensory proteins and their signaling networks. Semin Cell Dev Biol 92: 114–121

50. Podolec R, Ulm R (2018) Photoreceptor-mediated regulation of the COP1/SPA E3 ubiquitin ligase. Curr Opin Plant Biol 45: 18–25

51. Roman thank G, Rothenberg M, Lehman A, Ohme-Takagi M, Sinshi H, Klee H, Dolan L, Chang C, Roberts K, Quail PH, et al (1994) Meyerowitz for preprints of manuscripts or communication of results before publica-tion. also thank H. Klee for providing photographs of the tomato Nr mutant Phytochromes: Photosensory Perception and Signal Transduction.

52. Saijo Y, Sullivan JA, Wang H, Yang J, Shen Y, Rubio V, Ma L, Hoecker U, Deng XW (2003) The COP1-SPA1 interaction defines a critical step in phytochrome A-mediated regulation of HY5 activity. Genes Dev 17: 2642–2647

53. Shah J, Zeier J (2013) Long-distance communication and signal amplification in systemic acquired resistance. Front Plant Sci. doi: 10.3389/fpls.2013.00030

54. Singh D, Patil V, Kumar R, Gautam JK, Singh V, Nandi AK (2023) RSI1/FLD and its epigenetic target RRTF1 are essential for the retention of infection memory in Arabidopsis thaliana. Plant Journal 115: 662–677

55. Singh N, Giri MK, Chattopadhyay D (2025) Lighting the path: how light signaling regulates stomatal movement and plant immunity. J Exp Bot 76: 769–786

56. Singh N, Nandi AK (2022) AtOZF1 positively regulates JA signaling and SA-JA cross-talk in Arabidopsis thaliana. J Biosci. doi: 10.1007/s12038-021-00243-6

57. Singh N, Swain S, Singh A, Nandi AK (2018) AtOZF1 positively regulates defense against bacterial pathogens and NPR1-independent salicylic acid signaling. Molecular Plant-Microbe Interactions 31: 323–333

58. Singh V, Roy S, Giri MK, Chaturvedi R, Chowdhury Z, Shah J, Nandi AK (2013) Arabidopsis thaliana FLOWERING LOCUS D is required for systemic acquired resistance. Molecular Plant-Microbe Interactions 26: 1079–1088

59. Uquillas C, Letelier I, Blanco F, Jordana X, Holuigue L (2004) NPR1-Independent Activation of Immediate Early Salicylic Acid-Responsive Genes in Arabidopsis.

60. Velásquez AC, Castroverde CDM, He SY (2018) Plant–Pathogen Warfare under Changing Climate Conditions. Current Biology 28: R619–R634

61. De Wit PJGM (2007) How plants recognize pathogens and defend themselves. Cellular and Molecular Life Sciences 64: 2726–2732

62. Wu L, Yang HQ (2010) CRYPTOCHROME 1 is implicated in promoting R protein-mediated plant resistance to pseudomonas syringae in arabidopsis. Mol Plant 3: 539–548

63. Wu W, Chen L, Liang R, Huang S, Li X, Huang B, Luo H, Zhang M, Wang X, Zhu H (2024) The role of light in regulating plant growth, development and sugar metabolism: a review. Front Plant Sci. doi: 10.3389/fpls.2024.1507628

64. Xiang S, Wu S, Jing Y, Chen L, Yu D (2022) Phytochrome B regulates jasmonic acid-mediated defense response against Botrytis cinerea in Arabidopsis. Plant Divers 44: 109–115

65. Zavaliev R, Dong X (2024) NPR1, a key immune regulator for plant survival under biotic and abiotic stresses. Mol Cell 84: 131–141

66. Zeier J, Pink B, Mueller MJ, Berger S (2004) Light conditions influence specific defence responses in incompatible plant-pathogen interactions: Uncoupling systemic resistance from salicylic acid and PR-1 accumulation. Planta 219: 673–683

67. Zhou Y, Liu P, Tang Y, Liu J, Tang Y, Zhuang Y, Li X, Xu K, Zhou Z, Li J, et al (2024) NPR1 promotes blue light-induced plant photomorphogenesis by ubiquitinating and degrading PIF4. Proc Natl Acad Sci U S A. doi: 10.1073/pnas.2412755121

